# Varicose-projection astrocytes: a reactive phenotype associated with neuropathology

**DOI:** 10.1101/2025.03.17.643488

**Authors:** Caterina Ciani, Giulio Pistorio, Simona Giancola, Marika Mearelli, Francesca Emma Mongelli, Delaram Forouzeh, Luca Mio, Paula Ramos-Gonzalez, Carlos Matute, Ugne Kuliesiute, Sari Elena Dötterer, Gediminas Luksys, Saulius Rocka, Urte Neniskyte, Marya Ayub, Jean-Marie Graïc, Francesco Petrelli, Alexei Verkhratsky, Nunzio Iraci, Fabio Cavaliere, Carmen Falcone

## Abstract

Glial cells are fundamental for the pathophysiology of all neurological disorders. Astrocytes, the primary homeostatic cells of the central nervous system (CNS), exhibit species-specific characteristics, with human astrocytes specifically displaying unique structural and functional features. It is thus essential to investigate human-specific astrocytic responses to neuropathology using human-relevant models. Varicose projection (VP) astrocytes, traditionally considered specific to humans and apes, were suggested to reflect pathological burden, albeit direct evidence linking them to neurological diseases has been lacking. Here, we demonstrate for the first time that VP astrocytes are present in mice and tigers (*Panthera tigris*) and we provide evidence from four distinct human-based models that VP astrocytes are not a distinct physiological astrocyte subtype but rather a novel class of reactive astrocytes associated with neuropathology. Using human induced pluripotent stem cell (hiPSC)-derived astrocytes, mixed neural cultures, and cortical organoids, we showed that VP astrocytes are induced by pro-inflammatory cytokines interleukin-1***β*** (IL-1***β***) and tumor necrosis factor-***α*** (TNF-***α***) or LPS. Notably, cytokine withdrawal reverses the VP phenotype of astrocytes, indicating that it is a transient, inflammation-dependent state. We characterized the distinctive components of varicosities, including markers for extracellular vesicles, mitochondria, Golgi and endoplasmic reticulum components, suggesting roles in cellular stress responses and metabolic dysregulation. We further validated the pathological relevance of VP astrocytes by documenting their significant enrichment in postmortem brain samples from patients with several neurodegenerative diseases including as Alzheimer’s disease, Parkinson’s disease, and multiple sclerosis, as well as in surgical resections from patients with epilepsy due to hippocampal sclerosis or brain tumors, including previously unreported subcortical regions such as basal ganglia. Additionally, we identified a higher number of VP astrocytes also in mouse astrocytes upon treatment with pro-inflammatory cytokines, suggesting that formation of VP astrocytes is an evolutionarily conserved astrocytic response to neuroinflammation. Our findings point to VP astrocytes as a novel reactive astrocyte subtype closely linked to neuropathology, highlighting their potential as biomarkers and therapeutic targets in neurological diseases. This study lays the groundwork for future investigations into the mechanisms driving VP astrocyte formation and their broader implications in neuropathology.

## 1 Introduction

Understanding the mechanisms underlying astrocyte reactivity and its role in pathology remains a major challenge in neurobiology[1]. Astrocytes are essential regulators of brain homeostasis, supporting synaptic function, maintaining the blood-brain barrier, and modulating neuronal activity[2]. In pathological conditions, astrocytes undergo profound morphological and functional changes, a process known as reactive astrogliosis. Astrocyte reactivity is typically characterized by pronounced morphological alterations, including hypertrophy of the astrocyte soma, thickening and elongation of primary processes, upregulation of intermediate filaments such as GFAP, and the loss of their typical territorial (tiling) organization, sometimes accompanied by formation of overlapping processes[1, 3–5]. Reactive astrocytes can contribute to neuroprotection or, through loss of function, exacerbate neurodegeneration and disease progression, depending on the nature of the stimulus and the brain microenvironment[6, 7]. One prominent pathological context in which astrocyte reactivity plays a critical role is neuroinflammation, which encompasses a complex tissue response triggered by harmful inputs, such as trauma, disease, infections, and ischemia, accompanied with the production of pro-inflammatory molecules (including cytokines, small-molecule messengers, and chemokines)[8–11]. Inflammation of the nervous tissue translates to various consequences, such as synaptic malfunction, neuronal death, impaired neurogenesis, free radical depletion, cellular repair, and global gliosis[8, 9]. When astrocyte reactivity persists or is dysregulated, it may contribute to chronic neuropathological states[4, 8, 9, 11].

Astrocytes interact closely with microglia to enter an inflammatory state[6, 7, 11, 12]. In response to brain injury or toxic insults, microglia transform into a reactive state, secreting – among other factors – pro-inflammatory molecules such as tumor necrosis factor *α* (TNF-*α*), interleukin-1*α* (IL-1*α*), and the complement component subunit1q (C1q). These cytokines stimulate astrocytes, causing them to become reactive. This intricate crosstalk between astrocytes and microglia is fundamental for shaping the inflammatory landscape in the brain and determining the outcome of neuroinflammatory responses[7, 12, 13]. The consequences of astrocyte reactivity are widespread and heterogeneous, ranging from synaptic damage and reduced neuronal connectivity, to increased immune cell infiltration, which amplifies inflammation and disrupts classical astrocyte functions. These malfunctions were implicated in multiple neurological and psychiatric conditions[2, 7, 11, 13].

Despite extensive research in rodent models, the specific contribution of astrocytes to human pathology remains insufficiently understood. Astrocytes exhibit significant species-specific differences, with human astrocytes being larger, more structurally complex, and functionally distinct from those of rodents[14–17]. These differences often prevent the direct translation of findings from rodent models to human neurological diseases. Studying astrocytes in human-based models is essential for capturing their full range of responses to pathology, including their role in neuroinflammation.

Varicose-projection (VP) astrocytes were previously hypothesized to be primate-specific and to appear only under certain brain conditions, but their exact function and significance remain unclear[18, 19]. VP astrocytes present a distinctive morphology: their somata localized in deep cortical layers (V, VI, and white matter) sends long processes containing evenly distributed varicosities and spanning in all directions, they express GFAP and are devoid of classical astrocytic tiling patterns[14, 18]. VP astrocytes were originally identified in hominoid species (i.e., humans and other apes)[14, 18]; varicosities however were observed in other mammalian glial cells, including radial glial cells in the early postnatal primary visual cortex of cats (postnatal day 22)[20], in white matter astrocytes of neonatal pigs under hypoxia conditions[21], and – more recently – in the white matter of the ferret cerebrum[22]. However, cells displaying similar morphological structures described previously in literature were often not classified as or referred to explicitly as VP astrocytes.

The occurrence of VP astrocytes exhibits interindividual variation as they are detected in a few but not all representatives of the same species[18], suggesting that they are not a constitutive astrocyte subtype but instead only emerge under specific brain conditions[18]. Furthermore, VP astrocytes were not observed in neonatal or infant human brains[18], and they appear in the same cortical regions where interlaminar astrocytes display varicosities, reinforcing the hypothesis that they arise in response to specific pathological stimuli rather than being a physiological subtype[18]. The activation of the STAT3 pathway in reactive astrocytes, leading to protein degradation and ‘bead-like structures’ (similar to varicosities)[23, 24], supports the hypothesis that VP astrocytes represent a reactive rather than a physiological morphotype, induced by pathological conditions.

Here, we show for the first time that VP astrocytes are induced by pro-inflammatory cytokines *in vitro*, as observed in both pure human induced-pluripotent stem cell (hiPSC)-derived astrocytes and mixed (neuronal-astrocytic) cultures exposed to IL-1*β* and TNF-*α* stimulation. This induction is reversible upon cytokine withdrawal. Furthermore, we show that VP astrocytes are induced in 3D cerebral organoid models containing microglia, where formation of VP astrocytes was triggered by lipopolysaccharide (LPS). Our molecular characterization of VP astrocytes indicates the presence of markers for extracellular vesicles (EVs), components of mitochondria, Golgi complex and endoplasmic reticulum, suggesting their potential involvement in cellular communication, complex intracellular trafficking, metabolic regulation, and stress responses.

To establish the pathological relevance of VP astrocytes, we quantified their density in human postmortem brains from individuals with neurodegenerative disorders, including Alzheimer’s disease (AD), Parkinson’s disease (PD) and multiple sclerosis (MS), as well as in cortical surgical resections from patients with epilepsy due to hippocampal sclerosis or cancer. Density of VP astrocytes was significantly increased in these pathological conditions compared to controls, reinforcing their association with disease states. Additionally, we identified VP astrocytes in mouse embryonic stem cell (mESC)-derived astrocytes and in postmortem brains of tigers (*Panthera tigris*), suggesting that VP astrocyte formation is not exclusive to hominoids but may represent an evolutionary conserved astrocyte response to cellular stress and neuropathology at large.

In conclusion, our study demonstrates that VP astrocytes represent a reactive astrocyte phenotype emerging in response to pathological conditions, rather than a physiological astrocyte subtype. By elucidating their formation, molecular features, and presence in neurological diseases, this research increases our understanding of astrocyte reactivity in brain pathology and provides a foundation for exploring VP astrocytes as potential biomarkers or therapeutic targets in neuroinflammatory and neurodegenerative disorders.

## 2 Results

### 2.1 VP astrocytes are not hominoid-specific but are present across mammals

VP astrocytes were thought to be exclusive to humans and other hominoids, as they were identified only in primate brains. However, our observations reveal their presence in other mammalian species, including mouse (Figure 1 a-c), indicating that VP astrocytes are not restricted to primates. We also identified VP astrocytes in tiger brains (Panthera tigris, Figure 1d-e), further supporting their presence in gyrencephalic carnivores, as previously reported for ferret[22]. This suggests that VP astrocytes may represent a conserved astrocytic type across diverse mammalian species, rather than being exclusive to primates. Notably, similarly to previous observations in hominoids, VP astrocytes in other mammals are scattered and individual-specific (for example, VP astrocytes were present in 3 out of 5 samples of tigers), meaning that not all individuals from the same species exhibit them. This pattern further supports our hypothesis that VP astrocytes are not a distinct physiological subtype but rather emerge under specific brain conditions. Given this, we tested whether neuropathology, and neuroinflammation in particular, may drive VP astrocytes formation.

**Fig. 1.**
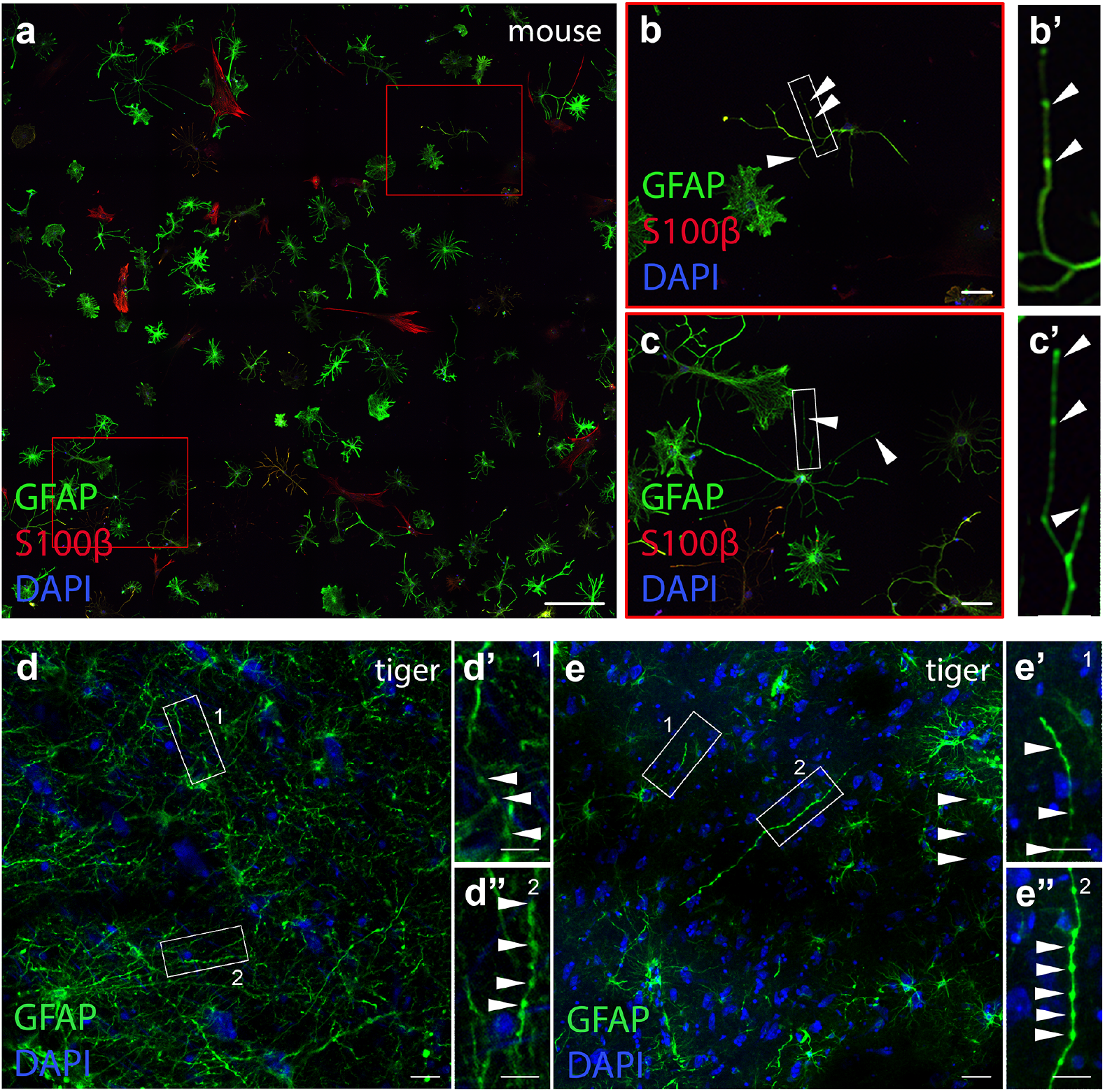
VP astrocytes are not hominoid specific but are found across mammals. (a-c) Representative immunofluorescence images of ESC-derived mouse astrocytes stained for GFAP (green), S100*β* (red), and DAPI (blue). (b,c) Higher magnification of the boxed regions in (a), highlighting varicosities (arrowheads). (b’,c’) Zoomed in images of the boxed regions in (b) and (c), respectively. (d-e”) GFAP immunostaining of postmortem prefrontal cortex tissue from a tiger (Panthera tigris), showing reactive astrocytes and VP Astrocytes. (d’–d”, e’-e”) Higher magnification of the boxed regions in (f, g), with arrowheads pointing to varicosities. Scale bars (a) = 100*µm*. Scale bars (b,c) = 25*µm*. Scale bar (e) = 25*µm*. Scale bars (d,e) = 50*µm*. Scale bars (d’-d”, e’-e”) = 25*µm*.

### 2.2 VP astrocytes are induced by pro-inflammatory cytokines in human astrocytes in vitro

To test our hypothesis, we differentiated human astrocytes from induced pluripotent stem cells (hIPSCs) in vitro, following an adapted protocol (see Methods). The astrocytic nature of the cultured cells was confirmed through immunostaining for canonical astrocytic markers, including S100*β*, GFAP, Aldh1l1, Aquaporin4 (AQP4), Kir4.1, EAAT1, Glutamine synthetase (GS), Sox9, and Vimentin (Supplementary Figure 1, Supplementary Figure 2). We treated these astrocytes with the proinflammatory cytokines IL-1*β* and TNF-*α*, either individually or in combination, at a concentration of 100ng/mL for each cytokine (see Methods). We evaluated the morphological and reactive changes in astrocytes upon these different treatments compared to the untreated control at different time points (1 hour, 24 hours, 72 hours, 5 days, and 7 days). At the 1-hour time point, we assessed astrocyte reactivity via immunofluorescence staining for nuclear factor *κ*B p65 (NF-*κ*B p65), a transcription factor activated during acute inflammation. In untreated astrocytes, the NF-*κ*B complex remained cytoplasmic (Supplementary Figure 3a-e’). Following cytokine treatment, NF-*κ*B p65 translocated to the nucleus, as shown by immunofluorescence (Supplementary Figure 3a-e’). We quantified the extent of nuclear translocation by calculating the nuclear-to-cytoplasmic intensity ratio, and found the strongest NF-*κ*B translocation under combined IL-1*β* and TNF-*α* treatment, with a three-fold increase compared to controls (Supplementary Figure 3a, IL-1*β* vs. Ctr: +69.6%, *p* = 0.0339; TNF-*α* vs Ctr: +240.9%, *p* < 0.0001; IL-1*β* /TNF-*α* vs. Ctr: +302.2%, *p* < 0.0001).

We further confirmed astrocyte reactivity by examining astrocyte morphology at later time points using GFAP immunostaining. By day 7, astrocytes exhibited significant morphological changes, including soma enlargement and increased number of processes. These changes were evident in IL-1*β*-treated cells (Supplementary Figure 3e’), TNF-*α*-treated cells (Supplementary Figure 3e”) and most prominently in the cells treated with both cytokines (Supplementary Figure 3e”‘), with the volume of GFAP+ astrocytes significantly higher compared to control (Volume of GFAP+ astrocytes after 7 days of treatment: Ctr= 13220.33 ± 1323.90 *µm*^3^, IL-1*β*+ TNF-*α* = 17433.89 ± 630.46 *µm*^3^, *p* < 0.04; Supplementary Figure 3d).

Next, we investigated the presence of VP astrocytes in these cultures through GFAP and S100*β* immunostaining at the same timepoints. We documented the appearance of VP astrocytes after 5 and 7 days of cytokine treatments in the IL-1*β* (Figure 2a,b,c-j’), TNF-*α* (Figure 2k-n’), and combined treatment conditions (Figure 2o-r’). In contrast, VP astrocytes were rare or absent in the control cultures at day 7, and were not observed in any condition prior to day 5 (Figure 2c-f). Quantification revealed up to a 45-fold increase in VP astrocytes in the combined treatment condition compared to controls, suggesting that the synergy between IL-1*β* and TNF-*α* was most effective in inducing varicosities (% VP astrocytes/total astrocytes in culture: Ctr = 0.07 ± 0.07%, IL-1*β* = 1.51 ± 0.60%, TNF-*α* = 1.07 ± 0.12%, IL-1*β*+ TNF-*α* = 3.20 ± 0.32%, with *p*[Ctr vs I vs. IL-1*β*]< 0.04, *p*[Ctr vs. TNF-*α*]< 0.001, *p*[Ctr vs. IL-1*β*+ TNF-*α*]< 0.0003; Figure 2b). Despite the increase, the overall frequency of VP astrocytes remained low, consistent with previous findings[18].

**Fig. 2.**
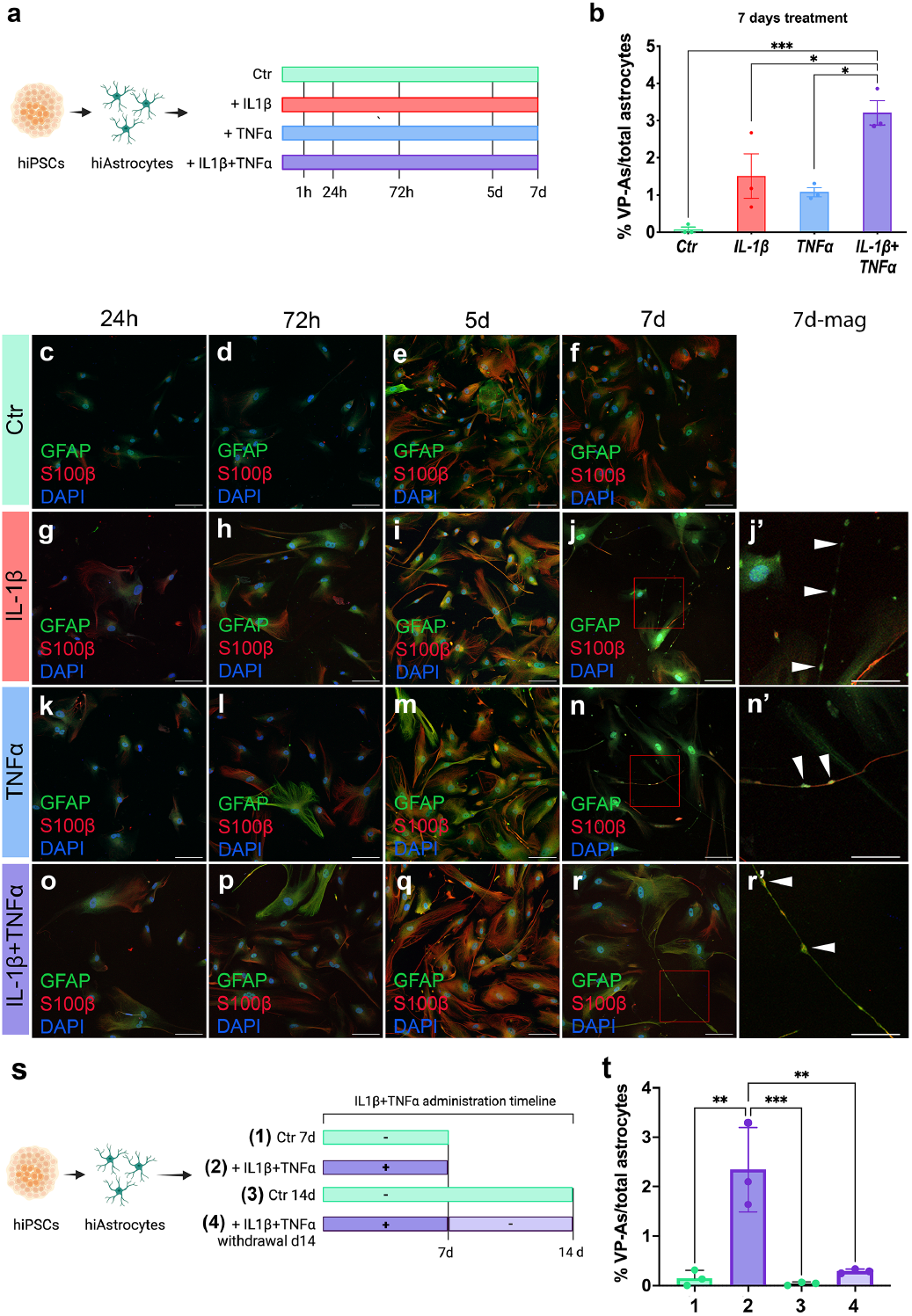
VP astrocytes are induced by pro-inflammatory cytokines in human astrocytes in vitro and their density increase is reversible one week after removing the cytokine exposure. (a) Schematic representation of the experimental protocol. Human iPSC-derived astrocytes were treated with interleukin-1*β* (IL-1*β*) and tumor necrosis factor-*α* (TNF-*α*) alone or in combination for up to 7 days. (b) Quantification of VP astrocyte density as a percentage of total astrocytes in culture across conditions. A significant increase in VP astrocytes is observed in cytokine-treated conditions, with the highest induction in the combined IL-1*β* + TNF-*α* treatment. Data are presented as mean ± s.e.m., statistical significance assessed by one-way ANOVA with Tukey’s multiple comparisons test (n = 3; * *p* < 0.05, ** *p* < 0.01, *** *p* < 0.001). (c–f) Representative immunofluorescence images of astrocytes under control (Ctr) conditions at (c) 24 hours, (d) 72 h, (e) 5 days, and (f) 7 d, showing the absence of VP astrocytes. h= hours; d= days. (g–j) Representative images of astrocytes treated with IL-1*β* at corresponding time points, demonstrating the progressive emergence of VP astrocytes, particularly at 5 d and 7 d. (j’) Higher magnification of the boxed region in (j), highlighting VP astrocytes. (k-n’, o-r’) Representative images of astrocytes treated with TNF-*α* (k-n’) and IL-1*β* + TNF-*α* (o-r’) at corresponding time points, showing an increased frequency of VP astrocytes, most notably in the combined condition. (n’) Higher magnification of the boxed region in (n), highlighting VP astrocytes. (r’) Higher magnification of the boxed region in (r), highlighting VP astrocytes. In all the fluorescent images, GFAP, S100*β*, and DAPI staining in green, red, and blue, respectively. White arrowheads point to varicosities. Scale bars (c-f, g-j, k-n, o-r) = 100 *µ*m. Scale bars (j’,n’,r’) = 50 *µ*m. (s) Schematic of the experimental timeline of the cytokine withdrawal experiment. Human iPSC-derived astrocytes were treated with IL-1*β* and TNF-*α* for 7 days, followed by cytokine withdrawal for an additional 7 days to assess reversibility. Four conditions were analyzed: (1) control at 7 days, (2) IL-1*β* + TNF-*α* treatment at 7 days, (3) control at 14 days, and (4) IL-1*β* + TNF-*α* withdrawal (rescue) at 14 days. (t) Quantification of VP astrocytes as a percentage of total astrocytes in each condition. Cytokine-treated astrocytes show a significant increase in VP astrocytes at 7 days, which is largely reversed upon cytokine withdrawal by day 14. Data are presented as mean ± s.e.m., statistical significance assessed by one-way ANOVA with Tukey’s multiple comparisons test (n = 3, * *p* < 0.05, ** *p* < 0.01, *** *p* < 0.001).

To further validate the link between pro-inflammatory stimulation and VP astrocytes, we repeated these experiments in mixed cultures containing mature astrocytes and neurons, thereby providing a more physiologically relevant environment. First, we confirmed astrocyte identity by immunostaining the cultures for the same astrocyte markers (Supplementary Figure 4, Supplementary Figure 5) and for neuronal marker MAP2 (Supplementary Figure 6a). We optimized cytokine concentrations for mixed culture to maintain cell viability, with 10 ng/mL of each cytokine identified as the optimal dose for inducing astrocyte reactivity (concentrations tested were: 1 ng/mL, 10 ng/mL, and 30 ng/mL of each cytokine for 7 days, Supplementary Figure 6). To assess the appearance of VP astrocytes in mixed cultures upon treatments with 10 ng/mL of IL-1*β*, 10 ng/mL of TNF-*α* or both combined, we immunostained the cells for GFAP and S100*β*, as before. We found a higher number of VP astrocytes in the cytokine-treated samples compared to control samples, where presence of varicosities was much lower, with a more pronounced increase in the combined treatment condition (Supplementary Figure 7).

These findings provide the first proof of principle that VP astrocytes are not a physiological subtype of astrocytes, but their presence might indeed be due to neuroinflammatory conditions.

### 2.3 VP astrocytes reverse after removing the cytokine exposure

To determine whether the presence of VP astrocytes in human astrocyte cultures is reversible, we conducted a cytokine withdrawal experiment. As in the experiment described in Figure 1, we treated hiPSC-derived astrocytes with a combination of proinflammatory cytokines IL-1*β* and TNF-*α*, at 100ng/mL each (Figure 2s). After one week, we confirmed the presence of VP astrocytes (Supplementary Figure 8b). We then removed the cytokines from the medium and cultured the astrocytes for an additional week. Notably, the number of VP astrocytes significantly decreased in the cytokine withdrawal condition (% VP astrocytes/total astrocytes in culture: Ctr at 7 days=0.15%, IL-1*β*+ TNF-*α* at 7 days=2.34± 0.49%, Ctr at 14 days= 0.04 ± 0.02%, IL-1*β*+ TNF-*α*-withdrawal at 14 days= 0.29± 0.02%, with *p*[IL-1*β*+ TNF-*α*(7days) vs. IL-1*β*+TNF-*α*(withdrawal-14 days)]< 0.007, *p*[IL-1*β*+ TNF-*α*(7days) vs. Ctr(7days)]= ns, *p*[Ctr(14days) vs. IL-1*β*+TNF-*α*(withdrawal-14 days)] < 0.0007, Figure 2t, Supplementary Figure 8). These findings indicate that VP astrocytes are a transient phenomenon, appearing only under sustained neuroinflammatory conditions. This reversibility highlights the dynamic nature of VP astrocytes, suggesting that interventions targeting inflammation could mitigate their formation.

### 2.4 VP astrocytes are induced by pro-inflammatory cytokines in mouse astrocytes in vitro

To determine whether VP astrocytes are an inflammation-associated feature unique to humans, we conducted a similar experiment using mouse embryonic stem cell (mESC)-derived astrocytes. First, we confirmed the astrocytic identity of the mouse cells by immunostaining for established markers, including GFAP, S100b, Aldh1l1, Kir4.1, EAAT1, GS, Sox9 and Vimentin (Supplementary Figure 9, Supplementary Figure 10).

Next, we exposed the mESC-derived mouse astrocytes to IL-1*β*, TNF-*α*, or a combination of the two cytokines, each at a concentration of 100ng/mL (Figure 3a). Remarkably, after 7 days of stimulation, we observed an increase of VP astrocytes in mouse astrocyte cultures in all three different conditions of stimulation, marking the first identification of VP astrocytes in this species (% VP astrocytes/total astrocytes in culture: Ctr= 0.60 ± 0.11%, IL-1*β*=2.18 ± 0.59%, TNF-*α*=1.080.1%, IL-1*β*+TNF-*α*= 1.10 ± 0.14%, with p[Ctr vs. IL-1*β*] < 0.05, p[Ctr vs. TNF-*α*] < 0.007, p[Ctr vs. IL-1*β*+ TNF-*α*] < 0.01. As indicated by white arrowheads (Figure 3b-e), VP astrocytes with structural characteristics similar to human VP astrocytes were present across all three treatment conditions.

**Fig. 3.**
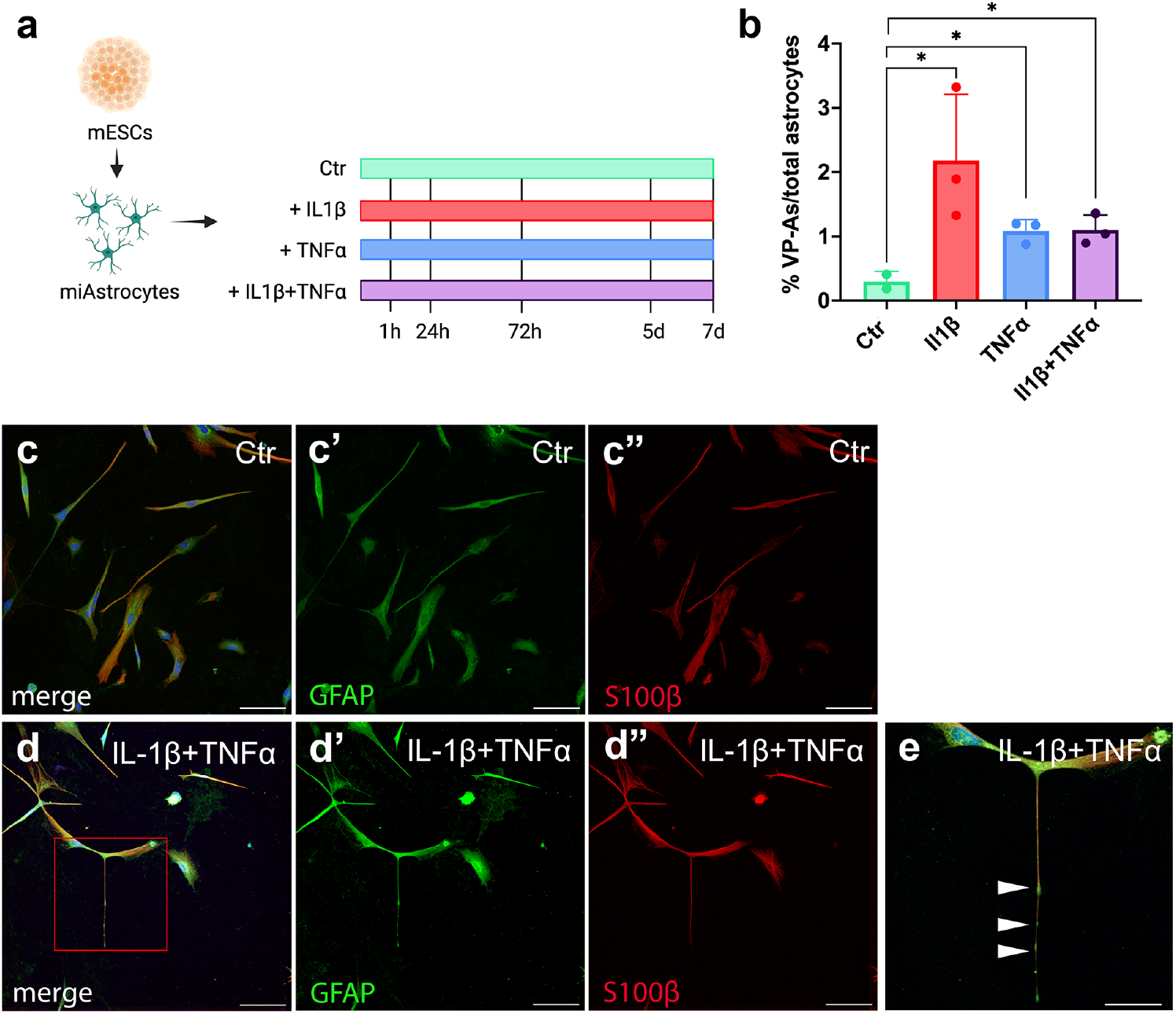
VP astrocytes are induced by pro-inflammatory cytokines in mouse astrocytes in vitro. (a) Schematic representation of the experimental timeline. Mouse embryonic stem cell (mESC)-derived astrocytes (miAstrocytes) were treated with IL-1*β*, TNF-*α*, or both cytokines for up to 7 days. (b) Quantification of VP astrocytes as a percentage of total astrocytes across conditions. A significant increase in VP astrocytes is observed following cytokine treatment compared to control. Data are presented as mean ± s.e.m., statistical significance assessed by one-way ANOVA with Tukey’s multiple comparisons test (n = 2,3,3,3, * *p* < 0.05, ** *p* < 0.01, *** *p* < 0.001). (c–e) Representative immunofluorescence images of mouse astrocytes stained for GFAP (green), S100*β* (red), and DAPI (blue) under control (c-c”) and IL-1*β* + TNF-*α*-treated (d-d”) conditions. (e) Higher magnification of the boxed regions in (d), highlighting varicosities (arrowheads).

These findings support the hypothesis that VP astrocytes are not a specific astrocyte subtype but rather a reactive phenotype induced by pro-inflammatory stimulation. This also suggests that VP astrocyte formation may represent a conserved astrocytic response to pathology across diverse mammalian species, rather than being exclusive to primates.

### 2.5 VP astrocytes are induced by pro-inflammatory cytokines in cerebral organoids

To further investigate the link between pro-inflammatory stimulation and VP astrocytes in a complex three-dimensional human model, we examined their presence in immunocompetent cerebral organoids (*i*.*e*., organoids with microglia). Cortical organoids were generated using an optimized protocol[25]. We differentiated organoids under gentle shaking for four months and added iPSC-derived microglia to two-month-old organoids (Figure 4a). To assess whether VP astrocytes could be induced by inflammatory stimuli, we treated four-month-old organoids with 10 ng/ml LPS for 24 hours, and performed GFAP immunofluorescence staining. Quantification revealed a significant increase in VP astrocyte density in LPS-treated organoids compared to controls (% VP astrocytes/total GFAP+ astrocytes: Ctr = 13.39 ± 3.65%; LPS = 60.66 ± 4.21%, *p* < 0.0001; Figure 4b-d”). These findings indicate that VP astrocytes emerge in response to inflammatory stimulation within a physiologically relevant human 3D system, reinforcing their association with neuroinflammatory conditions.

**Fig. 4.**
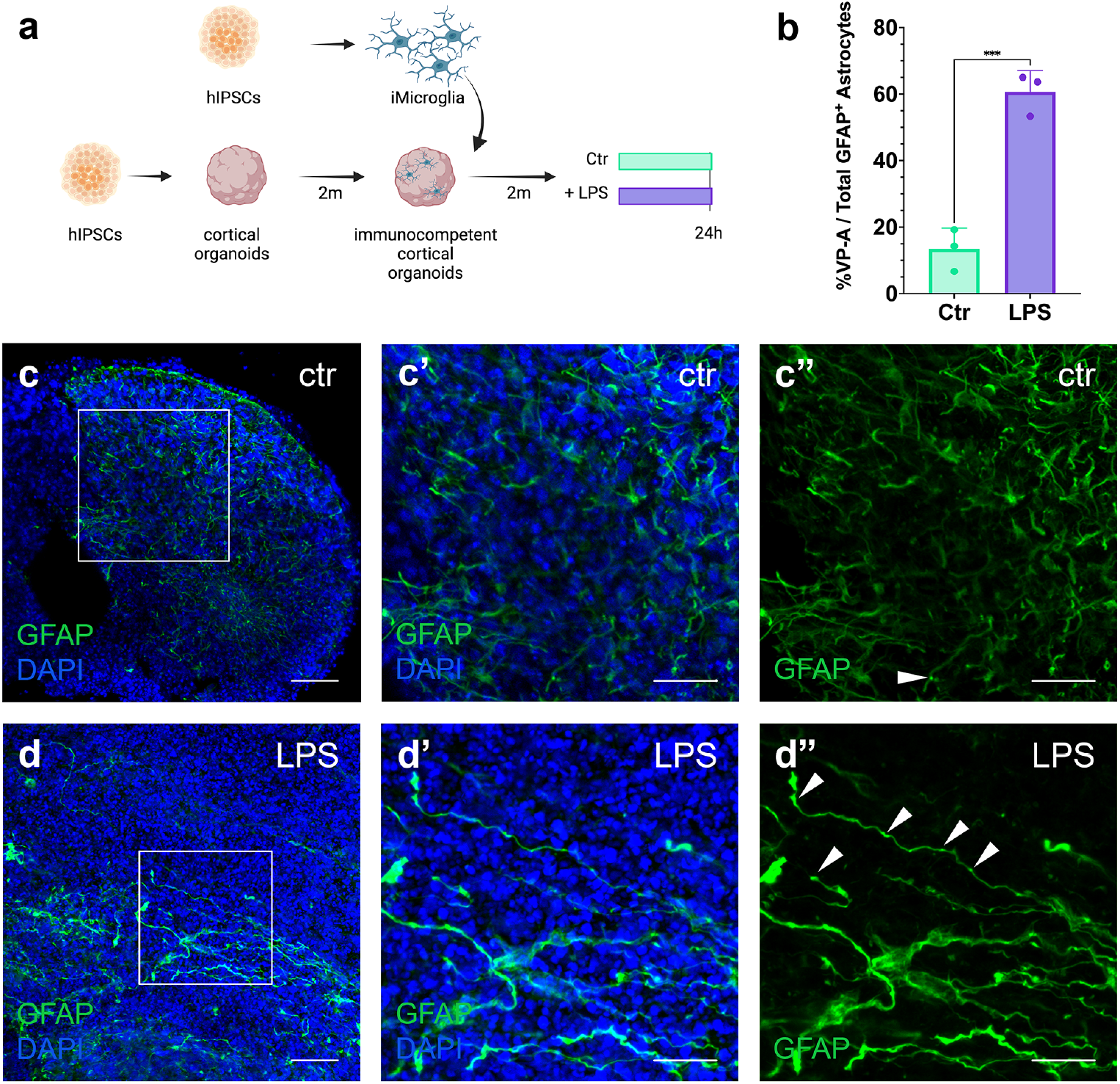
VP astrocyte density increases in immunocompetent cortical organoids after 24hour-exposure to LPS. (a) Schematic representation of the experimental timeline. hiPSCs were differentiated into cortical organoids for two months, then injected with induced microglia (iMicroglia) and cultured for an additional two months to generate immunocompetent cortical organoids. Organoids were treated with lipopolysaccharide (LPS) for 24 hours. (b) Quantification of VP astrocytes as a percentage of total GFAP+ astrocytes in control (Ctr) and LPS-treated conditions, showing a significant increase in VP astrocytes following LPS exposure. Data are presented as mean ± s.e.m., statistical significance assessed by two-tailed t-test (\*\**p* < 0.001). (c, d) Representative immunofluorescence images of GFAP+ astrocytes (green) and DAPI-stained nuclei (blue) in cortical organoids under control (c) and LPS-treated (d) conditions. (c’–c”) Higher magnification of GFAP+ astrocytes in control organoids, showing typical astrocyte morphology. (d’–d”) Higher magnification of GFAP+ astrocytes in LPS-treated organoids, with arrowheads indicating the presence of VP astrocytes. Scale bars (c,d) = 100 *µ*m; Scale bars (c’,c”,d’,d”) = 50 *µ*m.

### 2.6 Varicosities contain markers of extracellular vesicles and subcellular organelles

To elucidate the nature of the varicosities (the hallmark of VP astrocytes), we investigated whether they contain markers for EV components or specific cellular organelles, such as mitochondria, Golgi complex and endoplasmic reticulum (ER). We performed a treatment of hiPSC-derived astrocytes with 100 ng/ml each of IL-1*β* and TNF-*α* for 7 days, followed by immunofluorescence staining for various subcellular markers.

First, we stained astrocytes for GFAP (to label VP astrocytes) and EV markers, including Integrin 1, CD9, and CD63. Integrin *β*1, a multifunctional cell surface receptor involved in extracellular matrix interactions and EV uptake[26], was localized within the varicosities (Figure 5a-a”‘). Similarly, tetraspanin family proteins Cluster of differentiations 9 (CD9) and Cluster of differentiations 9 (CD63), key components of EV membranes, and involved in their biogenesis[27, 28], co-localized with varicosities (Figure 5b-b”‘ and 5c-c”‘, respectively).

**Fig. 5.**
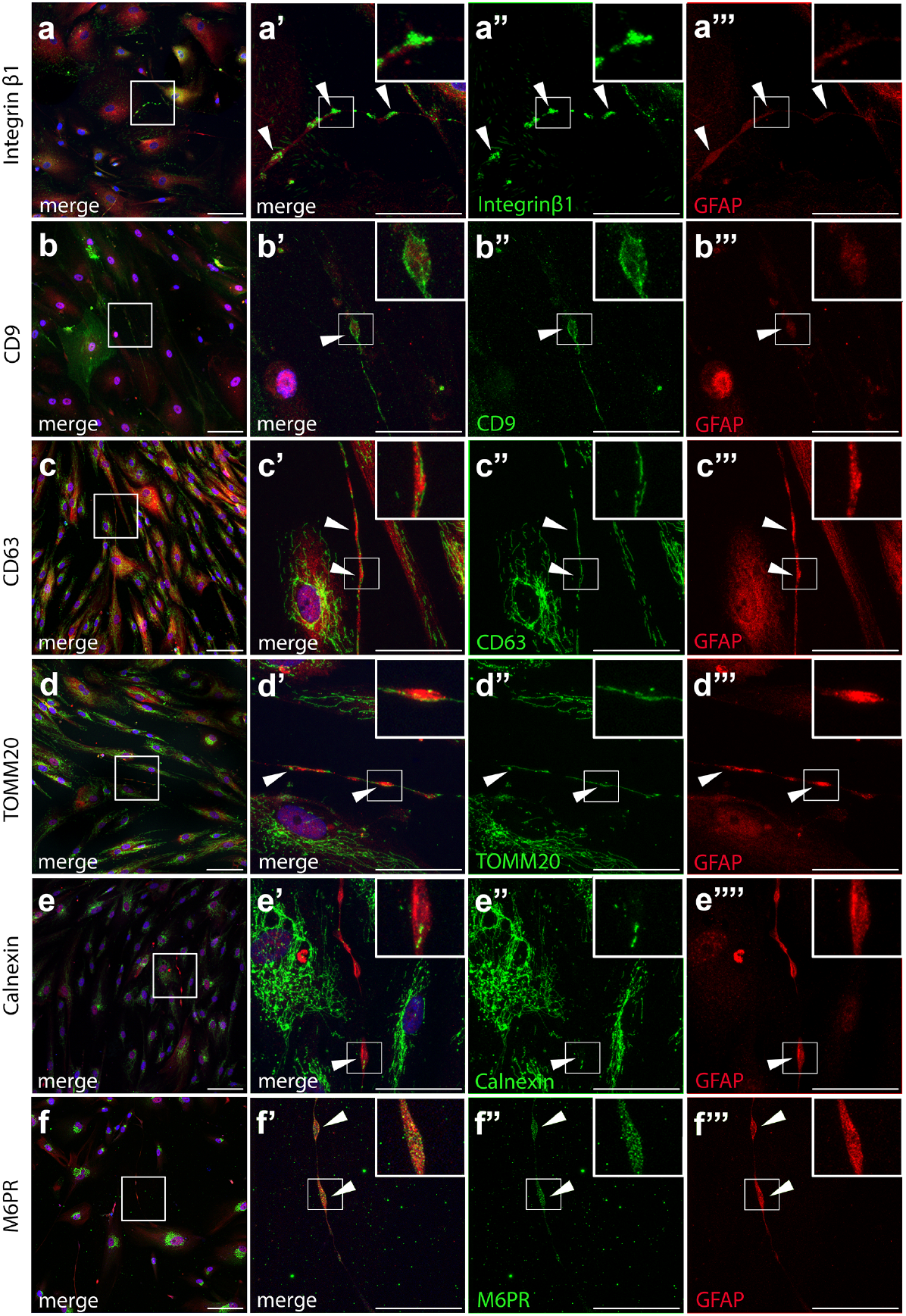
VP astrocytes contain extracellular vesicle markers and subcellular organelles. (a–a”‘) Representative immunofluorescence images of astrocytes stained for GFAP (red), Integrin *β*1 (green), and DAPI (blue). Integrin *β*1 co-localizes with varicosities (arrowheads). (a’-a”‘), Higher magnification of the boxed region in (a), highlighting varicosities. (b–b”‘) Representative immunofluorescence images of astrocytes stained for GFAP (red), CD9 (green), and DAPI (blue). CD9 co-localizes with varicosities (arrowheads) (b–b”‘), Higher magnification of the boxed region in (b), highlighting varicosities. (c–c”‘) Representative immunofluorescence images of astrocytes stained for GFAP (red), CD63 (green), and DAPI (blue). CD63 co-localizes with varicosities (arrowheads). (c–c”‘), Higher magnification of the boxed region in (c), highlighting varicosities. (d–d”‘) Representative immunofluorescence images of astrocytes stained for GFAP (red), TOMM20 (green), and DAPI (blue). TOMM20, a mitochondrial outer membrane protein, reveals fragmented mitochondria within varicosities (arrowheads). (d–d”‘), Higher magnification of the boxed region in (d), highlighting varicosities. (e–e”‘) Representative immunofluorescence images of astrocytes stained for GFAP (red), Calnexin (green), and DAPI (blue). Calnexin, an ER membrane protein involved in protein folding, is detected within varicosities (arrowheads). (e–e”‘), Higher magnification of the boxed region in (e), highlighting varicosities. (f–f”‘) Representative immunofluorescence images of astrocytes stained for GFAP (red), M6PR (green), and DAPI (blue). M6PR, a marker of the trans-Golgi network, is detected within varicosities (arrowheads). (f-f”‘) Higher magnification of the boxed region in (f), highlighting varicosities. Scale bars (a,b,c,d,e,f) = 100 *µ*m. Scale bars (a’-a”‘,b’-b”‘,c’-c”‘,d’-d”‘,’-e”‘,f’-f”‘) = 50 *µ*m.

Next, we assessed the presence of organelles. Staining for TOMM20, a mitochondrial outer membrane protein[29], revealed portions of mitochondria within varicosities (Figure 5d-d”‘), possibly indicative of a role in metabolic regulation during inflammation. Calnexin, an ER membrane protein involved in protein folding and quality control[30], was also present in VP astrocytes, as evidenced by co-localization with GFAP at the level of varicosities (Figure 5e-e”‘). Lastly, we examined the mannose-6-phosphate receptor (M6PR), a marker for the trans-Golgi network (TGN)[31], and identified its presence in varicosities (Figure 5f-f”‘).

This characterization demonstrates that VP astrocytes harbor a range of EV markers and intracellular organelles, suggesting their potential role in cellular communication and stress responses under inflammatory conditions.

### 2.7 VP astrocyte density increases in patients with neurodegenerative diseases and epilepsy

If VP astrocytes accumulate under pathological conditions, particularly in the presence of heightened neuroinflammation, their density should be increased in human neurological diseases. To investigate this, we examined postmortem human prefrontal cortex samples from patients with neurodegenerative disorders known for their inflammatory components[32], including Alzheimer’s disease (AD), Parkinson’s disease (PD), and multiple sclerosis (MS), and compared them to control samples from individuals without neurological disorders. Immunofluorescence staining for GFAP revealed a significant increase in VP astrocyte density in all three diseases. Specifically, when we calculated the density of VP astrocytes as GFAP+ VP astrocytes/Total n. of GFAP+ astrocytes, AD showed a 2.6-fold increase in VP astrocyte density (AD analysis: Ctr= 10.85 ± 0.13%, AD= 27.79 ± 5.63%, *p* < 0.01; Figure 6a,d,d’,e,e’), PD showed a 2.3-fold increase (PD analysis: Ctr= 10.85 ± 0.13%, PD= 24.62 ± 4.22%, *p* < 0.01, Figure 6b,f,f’); and MS showed a 3.7-fold increase (MS analysis: Ctr=10.85 ± 0.13%, MS= 40.69 ± 6.18%, *p* < 0.004, Figure 6c,g,g’), in respect to healthy controls. Incidentally, we also observed varicosities in interlaminar astrocytes (ILAs) in pathological samples across all three diseases (data not shown). Notably, control specimens contained some (but much fewer) VP astrocytes, likely due to the advanced age of the specimens.

**Fig. 6.**
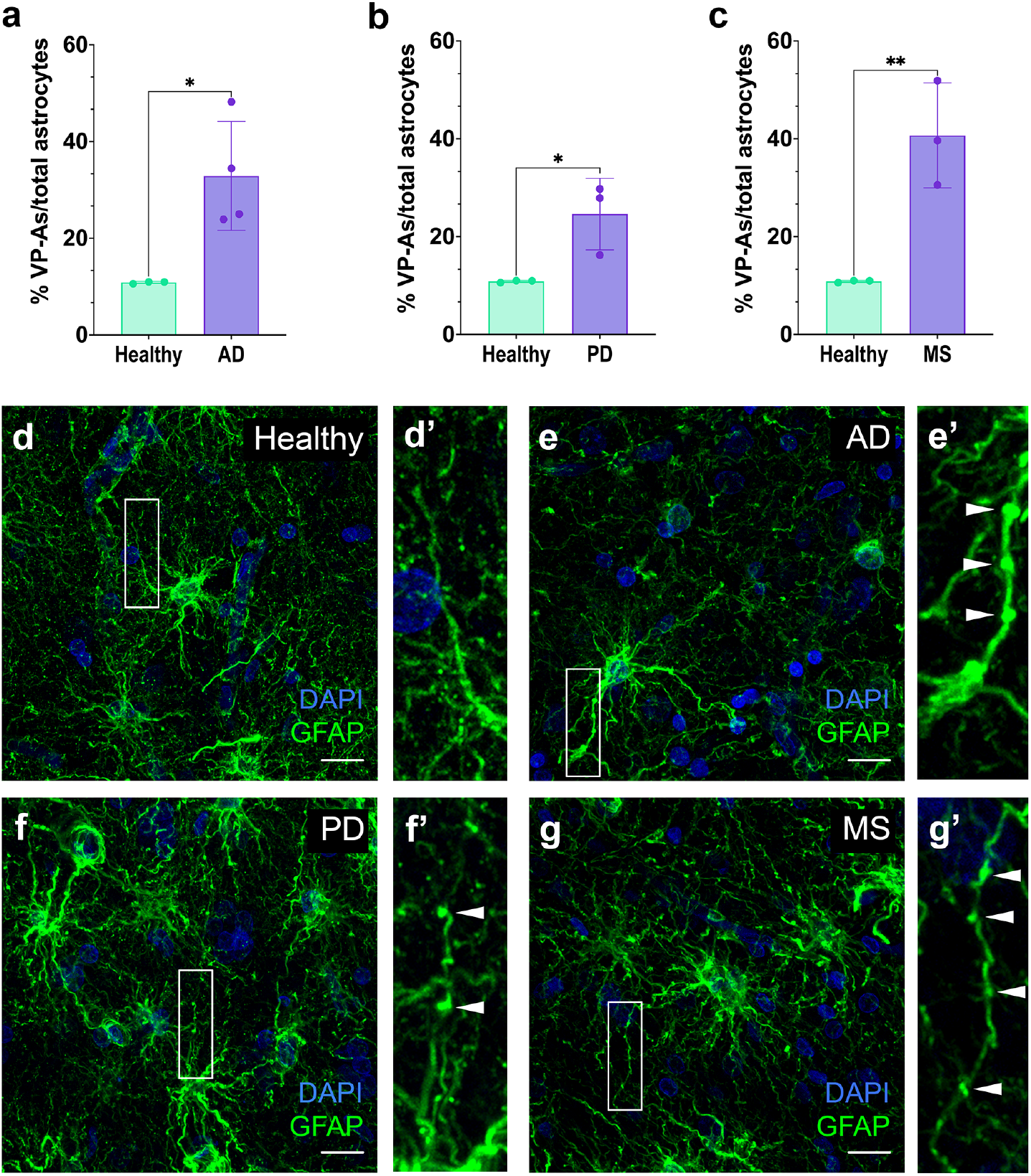
VP astrocyte density increases in the cerebral cortex in human neurodegenerative diseases. (a–c) Quantification of VP astrocytes as a percentage of total astrocytes in the prefrontal cortex of postmortem human brains from individuals with (a) Alzheimer’s disease (AD), (b) Parkinson’s disease (PD), and (c) multiple sclerosis (MS), compared to healthy controls. A significant increase in VP astrocytes is observed in all three pathological conditions. Data are presented as mean ± s.e.m., statistical significance assessed by two-tailed t-test (n = 3; * *p* < 0.05, ** *p* < 0.01, *** *p* < 0.001). (d–g) Representative immunofluorescence images of astrocytes stained with GFAP (green) and DAPI (blue) in healthy (d-d’) and diseased (e-e’) AD, (f-f-) PD, and (g-g’) MS conditions. GFAP (green) marks astrocytes, and DAPI (blue) stains nuclei. (d’,e’,f’, g’) Higher magnification of the boxed regions in (d), (e), (f), and (g), respectively, highlighting varicosities (arrowheads) in diseased conditions. Scale bars = 30 *µ*m.

In the samples from disease advanced stages, VP astrocytes were localized in all cortical layers and not only in the deepest ones, suggesting that this morphological change is a common astrocytic reaction and not something related to a single subpopulation of astrocytes. When the neurodegenerative disorder is at an advanced state, varicosities increase in number and size, consistently with the stressed state of the astrocytes (Figure 6e-g’). We found that VP astrocytes are detectable also by using another, well-known astrocyte marker S100*β* (Supplementary Figure 11).

To assess whether VP astrocytes are restricted to the cerebral cortex, we examined subcortical regions of postmortem PD brains, specifically the basal ganglia (BG) including the caudate nucleus, putamen and globus pallidus (BG-CPP) and subthalamic nucleus (BG-STN). VP astrocytes were present in these regions, with significantly increased densities in PD in both BG-CPP and BG-STN. Specifically we found approximately a 2 fold-increase in VP astrocyte density in BG-CPP (Density of VP astrocytes in BG-CPP calculated as GFAP+ VP astrocyte/Total n. of GFAP+ astrocytes: Ctr= 16.48 ± 1.11%, PD= 32.62 ± 0.92%, *p* < 0.0002, Figure 7a,c,d,g,h) and an approximate 3-fold-increase in BG-STN (Density of VP astrocytes in BG-STN calculated as GFAP+ VP astrocytes/Total n. of GFAP+ astrocytes: Ctr= 12.35 ± 1%, PD=38.96 ± 0.5%, *p* < 1.06*E*^*−*05^, Figure 7b,e,f,I,j).

**Fig. 7.**
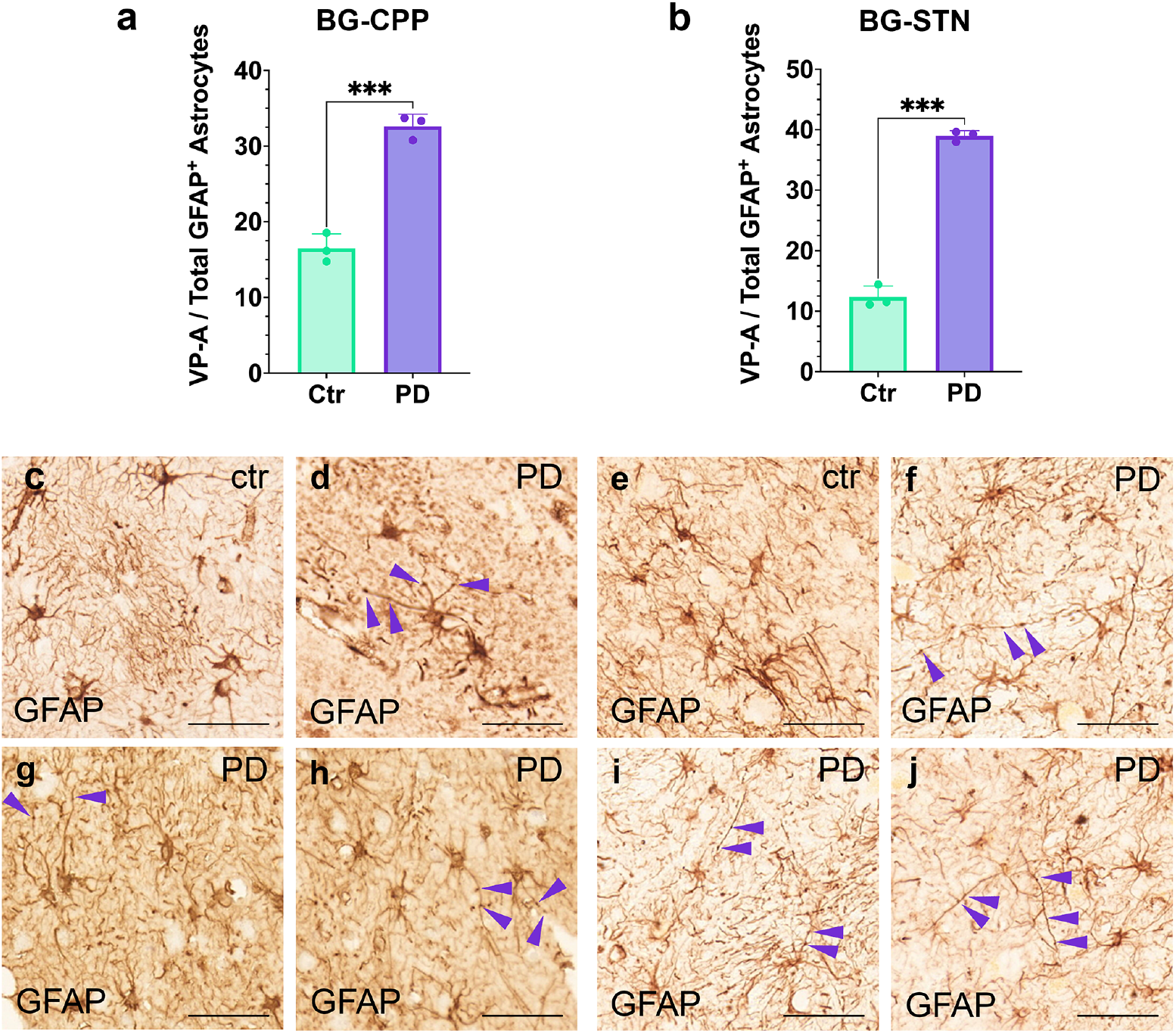
VP astrocyte density increases in subcortical regions of Parkinson’s disease. (a, b) Quantification of VP astrocytes as a percentage of total GFAP+ astrocytes in the (a) basal ganglia (BG-CPP) and (b) subthalamic nucleus (BG-STN) of postmortem human brains from Parkinson’s disease (PD) patients and healthy controls (Ctr). A significant increase in VP astrocytes is observed in PD in both regions. Data are presented as mean ± s.e.m., statistical significance assessed by two-tailed t-test (n = 3; * *p* < 0.05, ** *p* < 0.01, *** *p* < 0.001). (c–j) Representative immunohistochemistry images of GFAP+ astrocytes in control (c, e) and PD (d, g-j) samples from the BG-CPP (c–d, g–h) and BG-STN (e–f, i–j). VP astrocytes (arrowheads) are significantly more abundant in PD conditions compared to controls. Scale bars = 50 *µ*m.

We also investigated VP astrocyte density in surgical resection from patients with epilepsy caused by hippocampal sclerosis or brain tumors. VP astrocytes were significantly more frequent in patients with epilepsy due to hippocampal sclerosis, showing approximately a 2-fold increase in density (Density of VP astrocytes calculated as GFAP+ VP astrocytes/Total n. of GFAP+ astrocytes: Ctr= 0.69 ± 0.14%, Hippocampal sclerosis-Epilepsy = 1.33 ± 0.09%, *p* < 0.01, Figure 8a-c’); patients with epilepsy due to brain tumors showed similar density increase (Density of VP astrocytes calculated as GFAP+ VP astrocytes/Total n. of GFAP+ astrocytes: Ctr= 0.69 ± 0.14%, Tumor associated-Epilepsy = 1.59 ± 0.36%, *p* < 0.04, Figure 8a,d-d’).

**Fig. 8.**
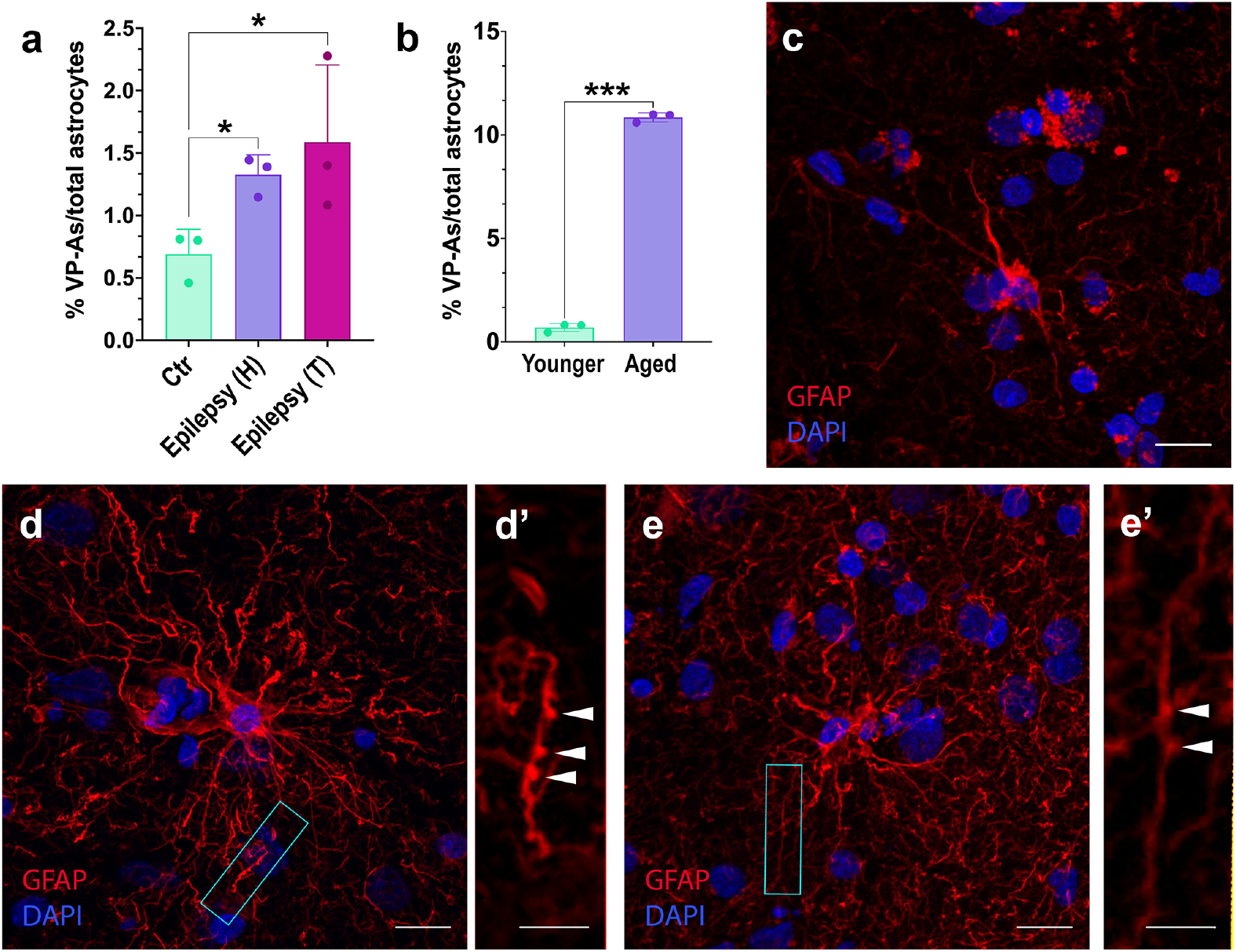
VP astrocytes are increased in epilepsy associated to hippocampal sclerosis and increase over time. (a) Quantification of VP astrocytes in surgical resections from patients with hippocampal sclerosis [Epilepsy (H)] and tumor-related epilepsy [Epilepsy (T)], showing a significant increase in both epilepsy due to hippocampal sclerosis and due to tumors, compared to control tissue. Data are presented as mean ±s.e.m., statistical significance assessed by two-tailed t-test (n = 3; * *p* < 0.05, ** *p* < 0.01, *** *p* < 0.001). (b) Quantification of VP astrocytes in young brains (from control surgical resections) vs. old brains (from control postmortem human brains). Data are presented as mean ± s.e.m., statistical significance assessed by two-tailed t-test (n = 3; * *p* < 0.05, ** *p* < 0.01, *** *p* < 0.001). (c–e’) Representative immunofluorescence images of GFAP+ astrocytes (red) and DAPI-stained nuclei (blue) in cortical tissue from controls (c), patients with epilepsy due to hippocampal sclerosis (d-d’) and patients with epilepsy due to brain tumor (e-e’). (d’, e’) Higher magnification of boxed regions in (d) and (e), respectively. showing prominent VP astrocytes (d’, e’, arrowheads). Scale bars (c,d,e) = 50 *µ*m. Scale bars (d’,e’) = 25 *µ*m.

These findings underscore that VP astrocytes are a hallmark of a subpopulation of reactive astrocytes in neuropathological conditions, with widespread occurrence across cortical and subcortical regions in both neurodegenerative and epileptic pathologies.

### 2.8 VP astrocytes accumulate with age

Previous studies have shown that VP astrocytes were not detected in neonatal or infant human brains[18, 19], suggesting that their presence may be associated with aging or pathological conditions. Given that aging is linked to the accumulation of various brain stressors and pathological burden, we investigated whether VP astrocyte density increases over time. To test this, we compared control brain tissues obtained from surgical resections of mid-age patients (age range = from 31 to 61 years old) with human postmortem brain samples from aged human controls (without diagnosed neurological disorders at the moment of death, age range = 77 to 81). Our analysis revealed a significant increase in VP astrocytes in older brains compared to younger ones, further supporting the idea that VP astrocytes may emerge in response to accumulated cellular stress or pathological processes associated with aging (Density of VP astrocytes calculated as GFAP+ VP astrocytes/Total n. of GFAP+ astrocytes: Younger= 0.69 ± 0.14%, Aged = 10.85 ± 0.16%, *p* < 10^*−*7^, Figure 8b).

## 3 Discussion

This study changes our understanding of VP astrocytes by establishing, for the first time, their presence in mice and tigers and their link to neuropathology and neuroinflammation, challenging the traditional view of varicosities as mere structural features of certain astrocytic subtypes. Our findings suggest that VP astrocytes are not a physiological astrocytic subtype but rather a reactive phenotype induced under specific pathological conditions. By employing multiple human-based models – including pure and mixed hiPSC-derived astrocyte cultures, 3D cortical organoids, human postmortem samples, and surgical resections – we provide robust evidence of their emergence in pathological conditions. This has significant implications for human neuropathology, given the species-specific differences in astrocyte properties and the unique features of human diseases[15, 32–35].

Our approach accounted for established pathways of astrocyte-microglia crosstalk[6, 7, 12, 36], using direct cytokine stimulation (TNF-*α*, IL-1*β*) in 2D cultures lacking microglia, and LPS-induced microglial activation in organoids[37], thus demonstrating that VP astrocytes arise from both direct and indirect inflammatory signaling.

Identifying VP astrocytes across multiple human pathologies underscores the potential of astrocyte varicosities as biomarkers for neuropathological processes, adding to previously used canonical markers such as GFAP upregulation and morphological hypertrophy. This raises intriguing questions about the molecular mechanisms that distinguish VP astrocytes from other reactive astrocytes, and whether VP astrocytes represent a specific astrocytic program or a pathway within the continuum of reactive states. Moreover, their increased presence in subcortical regions such as basal ganglia in PD (areas relevant to PD pathology) represents a completely novel finding, suggesting VP astrocytes respond adaptively according to regional vulnerability in the brain. Their presence across multiple cortical layers and subcortical regions suggests that VP astrocytes might play a role in propagating inflammation or exacerbating neuronal dysfunction. However, we cannot exclude a potential neuroprotective role. The overlap of varicosities between VP and interlaminar astrocytes in pathological states also warrants further investigation, as it hints at shared mechanisms or functions[18]. Additionally, the observed increase in VP astrocytes in aged human brains compared to younger samples suggests that their accumulation may be linked to age-related brain changes, potentially driven by chronic cellular stress or the gradual onset of pathological processes. This finding aligns with the idea that VP astrocytes are not a constitutive astrocyte subtype but rather emerge in response to accumulated physiological or pathological challenges over lifespan, reinforcing their potential role as markers of pathological risk.

Furthermore, we provide the first characterization of varicosities. We revealed the presence of EV markers, components of mitochondria, Golgi complex, and endoplasmic reticulum within varicosities, indicating potential roles in cellular communication, metabolic stress management, and intercellular signaling under pathological or inflammatory conditions, as previously reported for reactive astrocytes[26, 38, 39].

Our discovery that VP astrocytes are not restricted to hominoids[14, 18] but also appear in other mammals, like mice and tigers, challenges the prevailing view of their hominoid-specific nature. VP astrocyte formation may be therefore a conserved astrocytic response to neuropathology, underscoring their broader biological significance across species. This also raises important questions about the evolutionary pressures that shaped the reactive capabilities of astrocytes and whether similar structures exist in other vertebrates. One might question why VP astrocytes have not previously been described in other species. However, varicosities have indeed been reported in ferret astrocytes[22], in radial glial cells of early postnatal cats[20], and in neonatal pigs under hypoxia[21]. These earlier observations did not explicitly classify these structures as VP astrocytes. One explanation for their apparent scarcity in non-hominoid species could be that VP astrocytes accumulate progressively with aging, consistent with our hypothesis that their presence correlates with the accumulation of pathological burden over time.

Future work should focus on elucidating molecular mechanisms that regulate VP astrocyte formation, including pathways like NF-*κ*B [7, 40] and STAT3 signaling[1, 11, 41–44], their precise functional role in neuropathology, and whether they exacerbate neuronal damage or exert neuroprotective functions. The observed reversibility of VP astrocyte formation upon cytokine withdrawal indicates potential therapeutic avenues for modulating their emergence and pathological effects. The fact that VP astrocytes emerge in subcortical regions as well suggest that future studies should focus also on regional heterogeneity in VP astrocyte formation and its implications for disease-specific astrocytic dysfunctions. Finally, the potential interplay between VP astrocytes and other glial cells, such as microglia and oligodendrocytes, merits exploration, as glial crosstalk is increasingly recognized as a key driver of neuropathology.

This work aligns with and extends recent studies emphasizing the heterogeneity of astrocytic responses in neuropathology[1, 11]. Emerging evidence suggests that reactive astrocytes are not monolithic but exist on a spectrum influenced by context, region, and stimuli. VP astrocytes may represent a unique node on this spectrum, characterized by their distinct morphology and molecular composition, and open new avenues for translational studies in neuroinflammatory and neurodegenerative diseases.

## 4 Methods

### 4.1 Human induced pluripotent stem cell culture and differentiation into astrocytes

Human induced pluripotent stem cells (hiPSCs) were derived from normal human skin fibroblasts through episomal reprogramming methods. hiPSC line ASE-9209 (47y.o. female) was from Applied StemCell, Inc. hiPSCs were cultured and expanded in mTeSR™ Plus (STEMCELL Technologies, 05825) on Matrigel (Corning, 354277)-coated dishes, changing the medium every other day. hiPSCs were split using 1mL of ReLeSR (Stem cell technologies, 100-0484) for each well of the 6-well plate for 1 minute. Then 900 *µl* of ReLeSR was removed and cells were incubated for 7 minutes at 37^*°*^C. DMEM (Thermo Fisher, 31966047) was added to detach hiPSCs and collect them in a tube. Cells were centrifuged, then the supernatant was removed and resuspended in 1mL of mTeSR™ medium.

### 4.2 Human astrocyte cultures

hiPSCs were differentiated into astrocytes adapting a previously published protocol [45]. hiPSCs were split manually with ReLeSR (Stem cell technologies, 100-0484) to generate the embryoid bodies (*≈* day 5 of differentiation). Embryoid bodies were grown in suspension culture on a 6-well plate in DMEM/F-12 (Thermo Fisher, 11330032), with 1× N2 supplement (Invitrogen, 17502048) and Noggin (40 ng/ml, PeproTech, # 120-10C) for 3–5 days. Medium was changed daily. After 3-5 days, the Embryoid Bodies (EBs) were plated on Matrigel (Corning, 354277)-coated 6-well plate in Neural rosettes medium with DMEM/F12, 1× N2 supplement, and laminin (1 *µg/ml*, Invitrogen,2317-015) to form neural rosettes (=NPCs in form of neural rosettes) in 5-7 days. Medium was changed every other day. To differentiate immature astrocytes, the neural rosettes were mechanically dissociated and plated on a Matrigel-coated 6-well plate in astrocyte differentiation medium, consisting of DMEM/F12, 1× N2 supplement, 1× B27-RA supplement (Invitrogen, 12587010), BMP4 (10 ng/ml, Peprotech, 120-05ET), and FGF-basic (20 ng/ml, PeproTech, 100-18B). Medium was changed every other day.

### 4.3 Mixed culture model

hiPSCs were differentiated following a previously published protocol[46]. The colonies of hiPSCs were plated at 1× 105 cells per well on Matrigel-coated 6-well plate and fed daily with mTeSR™. After two days, differentiation, to generate OLIG2+ progenitors, was induced by adding neural induction medium with Retinoic acid (Sigma Aldrich, R265), SB431542 (Sigma Aldrich, 616461), and LDN193189 (Sigma Aldrich; SML0559) until day 8. On day 8, medium was switched to N2 medium until day 12 with 1× N2 supplement (Invitrogen, 17502048), changing medium daily. On day 12, cells were enzymatically dissociated using Accutase (Thermo Fisher, A1110501), breaking the monolayer into small clumps and transferring them into ultra-low attachment 6-well plate. N2B27 medium was added in each well with 1× N2 supplement and 1× B27 without vitamin A (Invitrogen, 12587010). Two-thirds medium change was performed every other day by transferring aggregates. On day 20, the medium was switched to PDGF-enriched medium using two-thirds medium change. These steps were repeated until day 30, to obtain the aggregation of OLIG2+ cells in spheres. On day 30, the round aggregates, with a diameter between 300 and 800 *µ*m and with a gold or brown center were picked with a p200 pipette, plated on a 6-well plate coated with poly-L-ornithine (Sigma Aldrich, P4957-50ML) and laminin (Sigma Aldrich, L2020-1MG), (20 spheres per well) and the medium was switched to glia medium. Medium was changed every other day replenishing two-thirds of the medium with fresh one until day 55. In this culture, are present mostly neurons and astrocytes, that can be maintained and expanded on a multi-well coated with poly-L-ornithine and laminin in glial medium.

### 4.4 Differentiation of mouse embryonic stem cells into astrocytes

The mouse embryonic stem cells (mESCs) were differentiated following a previously published protocol (Bien et al., 2016). The mESCs were dissociated using Accutase (Thermo Fisher, A1110501), and cultured on Matrigel (Corning, 354277) in ESCs medium before induction. On day 1, the medium was replaced by NE medium 1 comprising DMEM/F-12 supplemented with 1× B27 (Invitrogen, 12587010), 1× N2 (Invitrogen, 17502048), 1× NEAA (Euroclone, ECB3054D), 2-Mercaptoethanol (0.1 mM, Sigma Aldrich, M7154), Noggin (100 ng/ml, PeproTech,#120-10C), SB431542 (20 *µ*M, Sigma Aldrich; 616461), dorsomorphin (2 *µ*M, Sigma Aldrich, P5499-5MG), and CHIR99021 (3 *µ*M, Sigma Aldrich, SML1046-5MG). On day 4, neural rosettes-like colonies were dissociated with Accutase, and the medium was replaced with NE medium 2 consisting of DMEM/F12 supplemented with 1× N2, FGF2 (20 ng/ml, PeproTech/Immunotools, 100-18B), and glucose (1.6 g/L, Sigma Aldrich, G8270). The medium was maintained for an additional 4 days in ultra-low attachment plates to obtain neurospheres. On day 8, neurospheres were collected, dissociated with Accutase, and transferred to Matrigel-coated plates in NE media 3 with DMEM /F12, supplemented with 1× N2, 1× B27, Insulin (20 *µ*g /ml, Sigma Aldrich, 19278), glucose (1.6 g/L, Sigma Aldrich, G8270), EGF (20 ng/ml, Immunotools, 11343406), FGF2 (20 ng/ml, PeproTech/Immunotools, 100-18B). After 1 day, almost all cells dissociated from neurospheres being attached to the plate as NESCs, subsequently cultured in the NE media 3 to achieve cell expansion and differentiation in neurons and OPCs. On day 9, ESCs-derived NESCs were dissociated using Accutase and plated on a 6-well plate coated with Matrigel. Cells were grown for 5 days in OPC differentiation media with DMEM /F12, 1× B27, 1× N2, SAG supplements (0.4 *µ*M, Sigma Aldrich, 566661-500UG), FGF2 (20 ng/ml, PeproTech/Immunotools, 100-18B), PDGF-AA (20 ng/ml, Immunotools, 11343683). The resulting OPCs, grown and expanded in OPC media, are ready to be differentiated into astrocytes. All media were replaced every two days. Astrocytes were induced with Astro medium for 7 days with DMEM /F12, supplemented with 1× B27, 1× N2, and BMP4 (20ng/ml, PeproTech/Immunotools).

### 4.5 Cytokine treatments

The pure cultures of astrocytes were plated on a 12-multi well and treated with 100ng/mL of interleukin-1 beta (IL-1*β*) (Immunotools, 11340012), 100ng/mL of tumor necrosis factor-alpha (TNF-*α*) (PeproTech, 300-01A), or a combination of 100ng/mL + 100ng/mL of IL-1*β* and TNF-*α*. Controls were treated with 1× PBS, pH 7.4 (Thermo Fisher, 10010023). Starting concentration of 100ng/mL was chosen based on published protocols[47–49]. Astrocytes were treated for 1h, 24h, 72h, 5 and 7 days. The medium was not replaced throughout the treatments. The mixed culture was plated on a 12-well plate and treated with 4 different combined concentrations of cytokines for 7 days:

- 1ng/mL + 1ng/mL of IL-1*β* and TNF-*α*
- 10ng/mL + 10ng/mL of IL-1*β* and TNF-*α*
- 30ng/mL + 30ng/mL of IL-1*β* and TNF-*α*.
- 100ng/mL + 100ng/mL of IL-1*β* and TNF-*α*.

1× PBS, pH 7.4, was used as a treatment control. The medium was not replaced throughout the treatments. Mouse astrocytes were plated on a 12-well plate and treated with 100ng/mL of IL-1*β*, 100ng/mL of TNF-*α*, or a combination of 100ng/mL + 100ng/mL of IL-1*β* and TNF-*α*. 1× PBS, pH 7.4 (Thermo Fisher, 10010023), was used as a treatment control. The medium was not replaced throughout the treatments.

### 4.6 Human and animal tissues

The postmortem human samples were obtained from Netherlands Brain Bank. These specimens were fixed in formalin and paraffin-embedded. Prefrontal cortex was selected for the analysis. More detailed information regarding the human donors is provided in Table 1. The tissues were de-deparaffinized and embedded in OCT (Bio-Optica Milano S.p.A., #059801), following the protocol published in [50]. Then, 40*µ*m-thick sections were cut using a cryostat (Microm HM550) and stained by immunofluorescence (see below). Post-mortem paraffin-embedded brain samples included those from basal ganglia (BG) of Parkinson’s Disease (PD) patients, divided into two blocks per each individual: one block included the caudate nucleus, putamen and globus pallidus (BG-CPP) and another block included the subthalamic nucleus (BG-STN) (Table 1). These samples were obtained from the Biobank “Biobanc-Hospital Clinic-IDIBAPS” in Barcelona, Spain. The specimens of tigers (Panthera tigris) that died under human care were sent to the Department of Comparative Biomedicine and Food Science (BCA) at the University of Padua for routine postmortem examination. Brains were opportunistically gathered and immediately following extraction, were preserved by immersion in 4% phosphate-buffered formalin. Following death, autolysis begins to decompose biological tissues, including the brain. While in laboratory settings, smaller animals can be perfused with fixative right after euthanasia, maintaining minimal post-mortem intervals, larger animal brains collected during necropsies and strandings are stored in refrigerators at 4 ^*°*^C for several hours until the necropsy is conducted. This standard practice prevents damages caused by prolonged post-mortem intervals, thereby preserving tissue quality and staining results. The prefrontal cortex was sampled based on Musil and Olson [51]. We then embedded the tissues in OCT (Bio-Optica Milano S.p.A., #059801) as in [50], and cut 40*µ*m-thick sections at the cryostat (Microm HM550). Permission to use human brain tissue (from surgical resections) was obtained from Vilnius Regional Biomedical Research Ethics Committee (Approval No. 2020/2-1202-687). Human brain tissue specimens were obtained with the informed consent as requested by the Regional Ethics Committee (No. 2/2020 02 18). Human neocortical biopsy of tissue was sampled from either glioma tumor resection surgery (n = 3), using distant cortex without tumor infiltration (n = 3), or access cortex tissue from epilepsy surgery (n = 3) obtained during the resection of epileptic foci. Detailed information regarding the donors is provided in Table 1.

**Table 1.** List of human samples used in this study.

| ID | Sex | Age (y) | Brain region | Type of tissue | Pathology |
| --- | --- | --- | --- | --- | --- |
| 1 | M | 77 | PFC | Postmortem | Control |
| 2 | F | 81 | PFC | Postmortem | Control |
| 3 | M | 77 | PFC | Postmortem | Control |
| 4 | F | 79 | PFC | Postmortem | AD |
| 5 | F | 84 | PFC | Postmortem | AD |
| 6 | F | 88 | PFC | Postmortem | AD |
| 7 | F | 79 | PFC | Postmortem | PD |
| 8 | M | 83 | PFC | Postmortem | PD |
| 9 | F | 89 | PFC | Postmortem | PD |
| 10 | M | 78 | PFC | Postmortem | MS |
| 11 | F | 81 | PFC | Postmortem | MS |
| 12 | M | 78 | PFC | Postmortem | MS |
| 13 | F | 68 | STN | Postmortem | Control |
| 14 | M | 64 | STN | Postmortem | Control |
| 15 | F | 56 | STN | Postmortem | Control |
| 16 | M | 81 | STN | Postmortem | PD |
| 17 | F | 70 | STN | Postmortem | PD |
| 18 | M | 68 | STN | Postmortem | PD |
| 19 | F | 68 | BG | Postmortem | Control |
| 20 | M | 64 | BG | Postmortem | Control |
| 21 | F | 56 | BG | Postmortem | Control |
| 22 | M | 81 | BG | Postmortem | PD |
| 23 | F | 70 | BG | Postmortem | PD |
| 24 | M | 68 | BG | Postmortem | PD |
| 25 | M | 60 | Cortex | Surgical resection | Control |
| 26 | M | 61 | Cortex | Surgical resection | Control |
| 27 | M | 31 | Cortex | Surgical resection | Control |
| 28 | F | 28 | Cortex | Surgical resection | Ep/Hsc |
| 29 | F | 27 | Cortex | Surgical resection | Ep/Hsc |
| 30 | M | 47 | Cortex | Surgical resection | Ep/Hsc |
| 31 | F | 37 | Cortex | Surgical resection | Ep/T |
| 32 | M | 55 | Cortex | Surgical resection | Ep/T |
| 33 | M | 20 | Cortex | Surgical resection | Ep/T |

**Abbreviations:** M=male; F=female; y=years; PFC=Prefrontal cortex; STN: Subthalamic nucleus; BG= Basal ganglia; AD= Alzheimer's disease; PD= Parkinson' disease; MS=Multiple sclerosis; Ep/Hsc= Focal structural epilepsy with hippocampus sclerosis; Ep/T=Cortical epilepsy due to CNS tumour.

### 4.7 Cortical organoids

Brain organoids were generated from the standardized human iPSC cell line KOLF2.1J (The Jackson Laboratory) using an optimized protocol published by Lancaster et al.[25]. Briefly, each organoid was generated from 4.000 iPSCs using Aggrewell 800 (STEMCELL) using the commercial STEMDiff cerebral organoid kit (STEMCELL). Embryoid bodies generated were selected and, after 5 days in Aggrewell, differentiated (for 5 days) and matured under gentle shaking (80 rpm in Minitron shaker from Infors HT) for up to 4 months. iPSC-derived microglia from KOLF2.1J were introduced in 2 months old brain organoids. iPSCs were differentiated into hematopoietic progenitor cell (STEMdiff™ Hematopoietic Kit, STEMCELL) and differentiated two days in microglia before their inclusion by centrifugation (100 g, 3 minutes) in brain organoids (STEMdiff™ Microglia Differentiation Kit, STEM-CELL). Immunocompetent organoids generated were positive for the neuronal (SMI31, MAP2), astrocytic (GFAP, S100b), oligodendrocytic (Sox10, NogoA) and microglial (Iba1, CD68) markers. GFAP staining (DAKO 1.000) was performed by immunofluorescence in 4 months old organoids after 24 hours treatment with 10ng/ml of LPS.

### 4.8 Tissue processing

Postmortem tissues of prefrontal cortex from humans and tigers were fixed in formalin for several days, and subsequently incubated in 30% sucrose in PBS at 4^*°*^C for at least two days, allowing tissues to sink, which is essential for cryoprotection.

After cryoprotection, tissues were embedded in OCT medium, frozen on dry ice and stored at *−*80^*°*^C. OCT embedded tissue samples were cryosectioned (Microm HM550) into 40*µ*m-thick slices, collected as free-floating sections, and stored at *−* 20^*°*^C in anti-freeze medium until staining.

Postmortem tissues from the midbrain of PD patients and controls were fixed in formalin, paraffin-embedded and stored at room temperature until sectioning. Paraffin-embedded samples were sectioned with a vibratome into 40*µ*m-thick sections, and stored at 4^*°*^C until staining. Human surgical resection samples were placed in a cryomold, covered with an optimal cutting temperature (OCT) compound (Epredia™ Cryomatrix™ embedding resin, # 67-690-06, Fisher Scientific) and snap-frozen and stored at *−* 80^*°*^C until cryosectioning. OCT embedded tissue samples were cryosectioned (Cryotome^®^FE & FSE, A78910100, Thermo Fisher Scientific) into 20*µ*m slides and collected on SuperFrost Plus Adhesion slides (# 10149870, Thermo Scientific) and stored at *−*80^*°*^C until staining.

### 4.9 Immunofluorescence

For immunofluorescence on cell cultures, cells were fixed with paraformaldehyde (PFA) 4% in 1× PBS to perform immunofluorescence staining. We performed immunofluorescence first to characterize each model and then to evaluate the changes after treatment using GFAP and S100*β* markers. In the 12-well plate, cells were plated on 18mm coverslips coated with Matrigel for the monocultures or poly-L-ornithine and laminin for the mixed culture and then fixed with PFA 4%. To permeabilize the cells and reduce the a-specific binding of the antibodies, cells were incubated with Blocking Solution with 10% Donkey serum and 0,1% Triton for one hour. Later, cells were incubated with primary antibodies against the protein targets diluted in 1:5 of Blocking Solution and stored at 4^*°*^C overnight. The following day, cells were washed with 1× PBS three times for 10 minutes each. Then they were incubated with secondary antibodies conjugated with a fluorophore in 1:5 of Blocking Solution in 1× PBS for one hour. After that, cells were washed three times with 1× PBS. In case of the immunofluorescence staining for TOMM20 and GFAP was performed by incubating cells with secondary antibodies anti-rabbit 488 and anti-chicken biotinylated in Blocking Solution 1:5. Later, cells were washed three times for 10 minutes each in 1× PBS and then incubated with Streptavidin 594 (1:200, Invitrogen, S11227) for 30 minutes followed by three times washes in 1× PBS for 10 minutes each. Lastly, they were incubated with DAPI (1:1000, Merck-Sigma, 32670) in 1× PBS for 5 minutes at room temperature and then washed three times for 10 minutes each with 1× PBS. The coverslips were then mounted on a glass-slides sealed with Fluoromount-GTM (Invitrogen, 00-4958-02) letting them dry at room temperature in a dark box. We adapted a published immunofluorescence protocol for the staining experiments[50] for both cell cultures and postmortem human and tiger prefrontal cortex tissues. Postmortem human and tiger prefrontal cortex sections were immunostained free floating. A comprehensive list of primary and secondary antibodies used in this study is in Supplementary Table 1. Immunohistochemistry for GFAP expression on BG were performed in postmortem human brains from age-paired PD patients and non-PD patients. Brain sections were deparaffinized in xilol (5’) and decreased concentration of ethanol from 100% to 50% (5’ each) and washed in PB. After antigen retrieval (121^*°*^C), endogenous peroxidases were blocked for 30’ in 3% H2O2 in TBS and unspecific peptides in 4% BSA, 0.02% TritonX for 60 minutes. GFAP primary antibody (Dako, ref. Z0334) diluted 1:1,000 was incubated overnight in blocking solution (4% BSA, 0.02% TritonX) at 4^*°*^C. After three washes in TBS with 0.01% Triton, samples were incubated in biotinylated secondary antibody (mouse anti-rabbit 1:400) for two hours. After three washes in TBST, immunohistochemistry was developed in DAB (3,3’
s-diaminobenzidine tetrahydrochloride) substrate, following the company instruction (ThermoFisher). After, we performed dehydration in increasing concentration of ethanol (from 50% to 100% plus Xilene, each 5’). Immunofluorescence on human surgical resection samples was performed as follows. Sections were fixed in 4 % PFA in PBS for 15 min at room temperature, following with permeabilization in 1% Triton-X in PBS for 15 min and blocking in blocking buffer (10% normal goat serum, 2% bovine serum albumin in PBS) for 2 h. Sections were incubated with primary antibodies anti-GFAP (#ab4674, Abcam), anti-Iba1 (# 019-19741, FUJIFILM Wako Shibayagi) and anti-NeuN (#MAB377, Millipore) in antibody diluent buffer (2% normal goat serum in PBS) overnight at 4^*°*^C, visualized by secondary antibodies (Goat anti-Chicken IgY (H+L) Cross-Adsorbed Secondary Antibody Alexa Fluor™ Plus 594 (#A32759, Invitrogen), Goat anti-Mouse IgG (H+L) Cross-Adsorbed Secondary Antibody Alexa Fluor™ 488 (#A11001, Invitrogen) and Goat anti-Rabbit IgG (H+L) Highly Cross-Adsorbed Secondary Antibody, Alexa Fluor Plus 647 (#A32733, Invitrogen) in antibody diluent buffer for 2 h at room temperature. Immunolabeled sections were counterstained with DAPI (1 *µ*g/ml), washed and mounted with Mowiol. All antibodies used for this study are listed in the Supplementary Table 1.

### 4.10 Image acquisition and processing

The images from the immunofluorescence experiments on cell cultures, postmortem human prefrontal cortex, postmortem tiger prefrontal cortex and human surgical resection samples were acquired using a confocal microscope Nikon A1/R with different objectives (10X, 20X and 40X, SISSA microscope facility) based on the analysis we performed. We acquired every image using three or four lasers with different wavelengths: FITC (488 nm), TRITC (564 nm), Cy5 (647 nm) and DAPI (408 nm). Images were processed with “Fiji/ImageJ” software, merging the channels, and producing a Maximum Z-stack-projection. Images from postmortem human PD samples (BG-CPP and BG-STN) were taken in automated digital slide scanner Panoramic MIDI II by 3DHistech at the Achucarro’s Image Facility. NF-*κ*B translocation and astrocyte volume quantification To quantify the NF-*κ*B translocation and GFAP+ astrocyte volume we treated astrocytes for one hour, and then we acquired three images for each coverslip (3 coverslips for each treatment condition, n=3). Images were processed using “VolocityTM” software considering the intensity in the cytoplasm and the nucleus of each astrocyte and then we calculated the ratio of nuclei intensity over cytoplasm intensity to verify if the translocation had occurred. VP astrocytes quantification To characterize the varicose-projection astrocyte population, after treatment, we acquired large images for the 1-week treated immature astrocytes (n=3). We selected 3 points for each coverslip in the respective 3 spatial coordinates centering a varicose projection. For each point it was acquired an area of 4 × 4 fields with 20x magnification; this resulted in three large images for each coverslip, nine images for each condition at one week (Ctr, IL-1*β*, TNF-*α*, and IL-1*β*-TNF-*α*) with 20*µ*m of Z-stack with 19 steps, each of 1*µ*m. Images were processed into “Fiji/ImageJ” software, merging the channels, producing a Maximum Z-stack-projection, and exporting them as a Tiff file for the count. Varicosities were counted using the “Cell Counter” plugin in Fiji/ImageJ. We identified varicose projections along their corresponding originating cells. We considered only varicose projections when they exhibited more than two bulbous structures, discernible processes, and positive staining for either GFAP or S100*β*. The total number of cells in each large image was calculated using “VolocityTM” software with DAPI staining. Subsequently, we measured the percentage of VP Astrocytes over the total number of nuclei in the large images.

### 4.11 Statistical analysis

For NF-*κ*B translocation and VP astrocyte counts, data were obtained from at least three independent biological replicates within the same experiment. Prior to statistical analysis, data were tested for normality using the Shapiro-Wilk test. The values were then organized and exported into GraphPad Prism 8 software for statistical analysis. Depending on the comparison, results were analyzed using one-way analysis of variance (ANOVA) followed by Turkey’s multiple comparisons test or a two-tailed Student’s t-test. Statistical significance was set at *p* < 0.05 (* *p* < 0.05, ** *p ≤* 0.01, *** *p* < 0.001), and data are presented as mean ± standard error of the mean (SEM).

## Authors’ contribution

C.C. performed the experiments on postmortem human samples, coordinated the project and co-wrote the manuscript. G.P., S.G., M.M. and L.M. performed all the experiments with hiPSC-derived astrocytes and with mESC-derived astrocytes. F.M. performed the experiments and the analysis on postmortem human samples. D.F. performed the analysis from the human surgical resection samples. P.R.G., C.M, and F.C. performed the experiments on postmortem human samples (BG in PD), and cortical organoids. U.K. sample handling and preparation, immunofluorescent staining optimization and immunofluorescent staining for the surgical resections from human patients with epilepsy. S.E.D. performed the sample sectioning and immunofluorescent staining on the surgical resections from human patients with epilepsy. G.L. and S.R. provided the surgically resected human brain tissue. U.N. took care of conceptualization and funding acquisition for the results from the surgical resections from human patients with epilepsy. M.A. provided technical assistance for results on mouse cell experiments. J.M.G. provided the samples of the tiger and provided assistance with anatomical dissection of such samples. F.P. contributed to the conceptualization, analysis and interpretation of experiments for varicosity characterization, and to the writing of the manuscript. A.V. contributed to the interpretation of the data throughout the manuscript. N.I. provided the antibodies for EV markers, helped with the conceptualization of experiments for varicosity characterization and with the writing of the manuscript. C.F. conceptualized and directed the study, co-analyzed the results, made the figures and wrote the manuscript.

## Conflict of interest

The authors declare no conflict of interest.

## Acknowledgements

CF is grateful to the Human Technopole Early Career Fellowship that supported the bulk of this work. UN received funding from the European Regional Development Fund under grant agreement number 01.2.2-CPVA-V-716-01-0001 with the Central Project Management Agency (CPVA), Lithuania, and the Research Council of Lithuania under the Programme “University Excellence Initiatives” of the Ministry of Education, Science and Sports of the Republic of Lithuania (Measure No. 12-001-01-01-01 “Improving the Research and Study Environment”), project No. S-A-UEI-23-10. U.K. received funding from Vilnius University Science Advancement Fund, project No. MSF-JM-31/2022. PRG was awarded by the IKUR2030 program from the Basque Government. FC received funding from the Spanish Ministry of Science and Innovation (PGC2023. PID2023-146826OB-I00). We are grateful to Prof. Chet C. Sherwood, Dr. Matthew Broadhead and Dr. Fabio Anza for their insightful feedback and critical review of the manuscript, which greatly contributed to improving its clarity and scientific rigor.

## Notes

### Competing Interest Statement

The authors have declared no competing interest.

