## SupplementalMaterial for "Varicose-projection astrocytes: a reactive phenotype associated with neuropathology"


In this supplemental material we provide the following additional Figures in support of our Main Text:


- Supplemental Figure 1 – **Characterization of hiPSC-derived astrocytes (1)**
- Supplemental Figure 2 - **Characterization of hiPSC-derived astrocytes (2)**
- Supplemental Figure 3 - **Cytokine treatments and reactivity of hiPSC-derived astrocytes**
- Supplemental Figure 4 - **Characterization of hiPSC-derived astrocytes in mixed cultures (1)**
- Supplemental Figure 5 - **Characterization of hiPSC-derived astrocytes in mixed cultures(2)**
- Supplemental Figure 6 - **Cytokine treatments and reactivity of hiPSC-derived astroytes in mixed culture**
- Supplemental Figure 7 - **Varicose-projection astrocytes in treated human astrocytes in mixed culture**
- Supplemental Figure 8 - **VP astrocyte density increase is reversible one week after removing the cytokine exposure**
- Supplemental Figure 9 - **Characterization of ESC-derived mouse astrocytes (1)**
- Supplemental Figure 10 - **Characterization of ESC-derived mouse astrocytes (2)**
- Supplemental Figure 11 - **VP astrocytes express both GFAP and S100$\beta$**

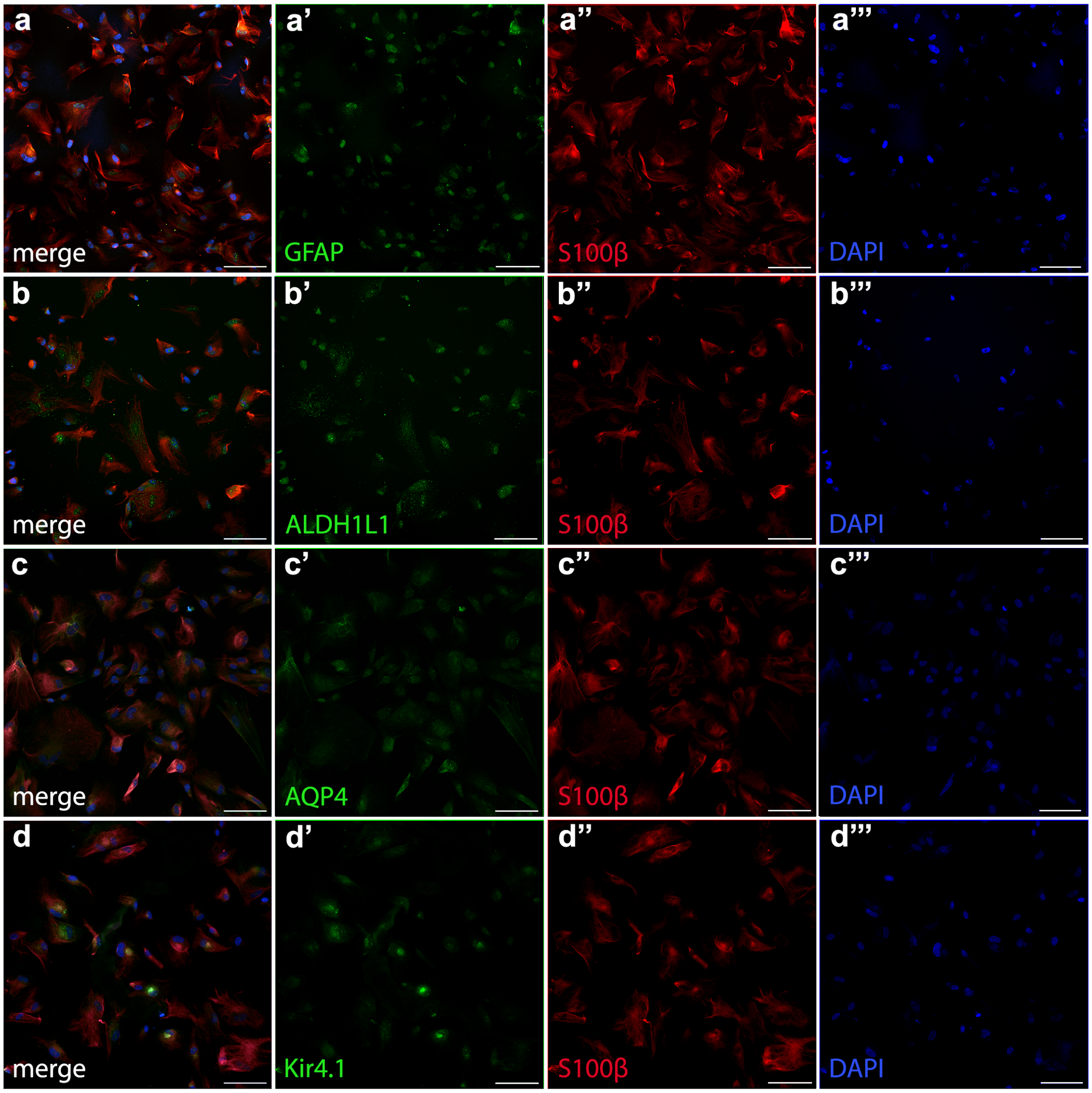


**Suppl. Fig. 1**: **Characterization of hiPSC-derived astrocytes (1).** (a–d) Representative immunofluorescence images showing the expression of key astrocytic markers in human induced pluripotent stem cell (hiPSC)-derived immature astrocytes. (a) GFAP (green), (b) ALDH1L1 (green), (c) AQP4 (green), and (d) Kir4.1 (green) are co-stained with the astrocytic marker S100$\beta$ (red) and nuclear marker DAPI (blue). Individual channels highlighting the expression pattern of GFAP (a'), ALDH1L1 (b'), AQP4 (c'), and Kir4.1 (d') in astrocytes. GFAP is predominantly localized in the nuclei and perinuclear region, ALDH1L1 is distributed in the astrocytic cytoplasm, AQP4 exhibits nuclear expression with lower levels in the astrocytic membrane, and Kir4.1 is enriched in the nuclear region. (a",b",c",d") S100$\beta$ (red) effectively delineates the morphology of hiPSC-derived astrocytes. (a"',b"',c"',d"') DAPI (blue) stains cell nuclei. Scale bars = 100 $\mu$m. These findings confirm the astrocytic identity of hiPSC-derived astrocytes based on the expression of multiple astrocytic markers.

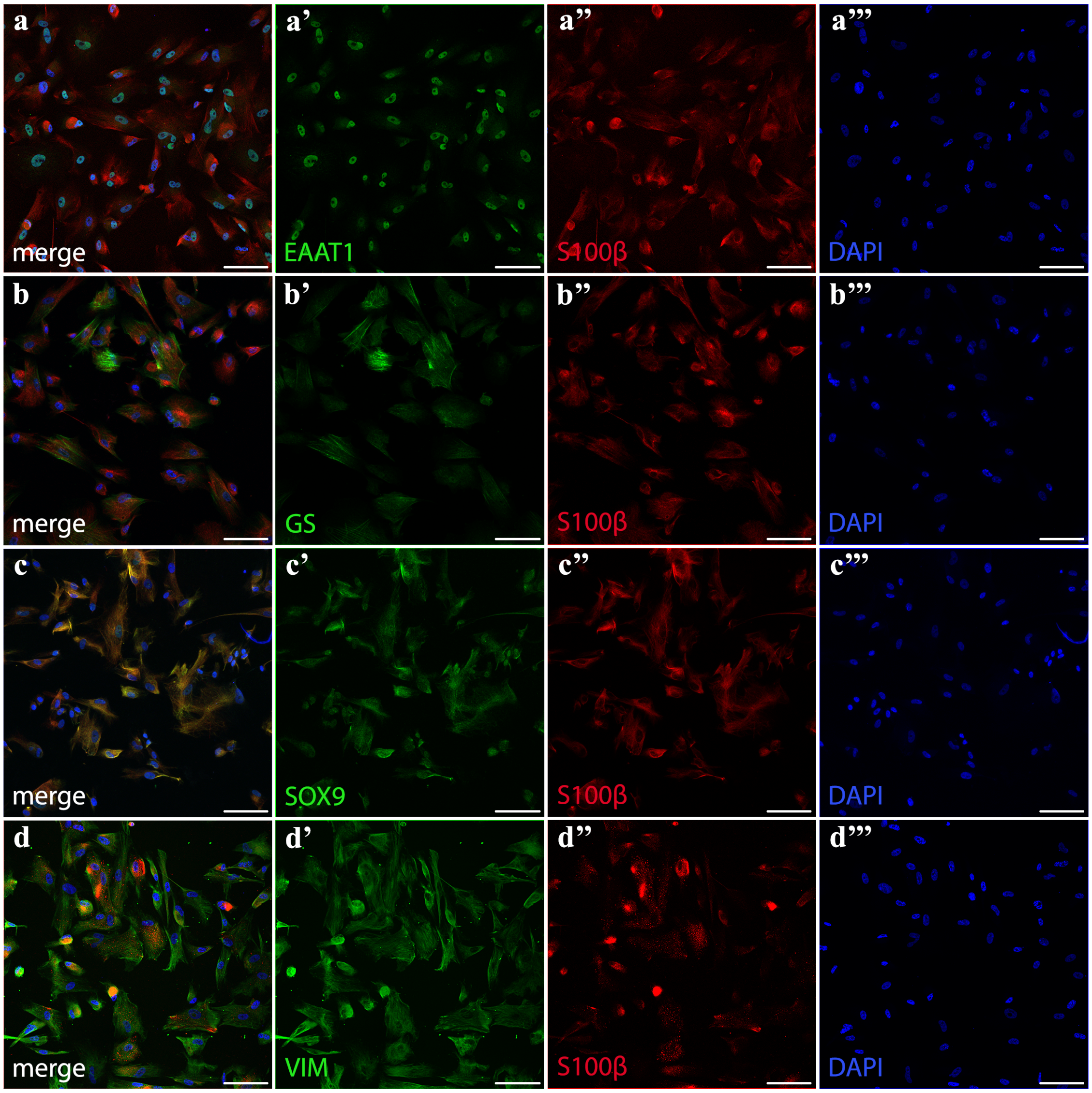


**Suppl. Fig. 2**: **Characterization of hiPSC-derived astrocytes (2).** (a–d) Representative immunofluorescence images showing the expression of additional astrocytic markers in human induced pluripotent stem cell (hiPSC)-derived immature astrocytes. (a) EAAT1 (green), (b) GS (green), (c) SOX9 (green), and (d) VIM (green) are co-stained with the astrocytic marker S100$\beta$ (red) and nuclear marker DAPI (blue). Individual channels highlighting the expression pattern of EAAT1 (a'), GS (b'), SOX9 (c'), and VIM (d') in astrocytes. EAAT1 is predominantly localized in the nucleus with faint expression in cell processes, GS is strongly expressed in the cytoplasm, SOX9 is present in both the nucleus and cytoplasm, and VIM outlines the morphology of immature astrocytes. (a",b",c",d") S100$\beta$ (red) effectively delineates the morphology of hiPSC-derived astrocytes. (a"',b"',c"',d"') DAPI (blue) stains cell nuclei. Scale bars = 100 $\mu$m. These findings further confirm the astrocytic identity of hiPSC-derived astrocytes through the expression of multiple astrocytic markers.

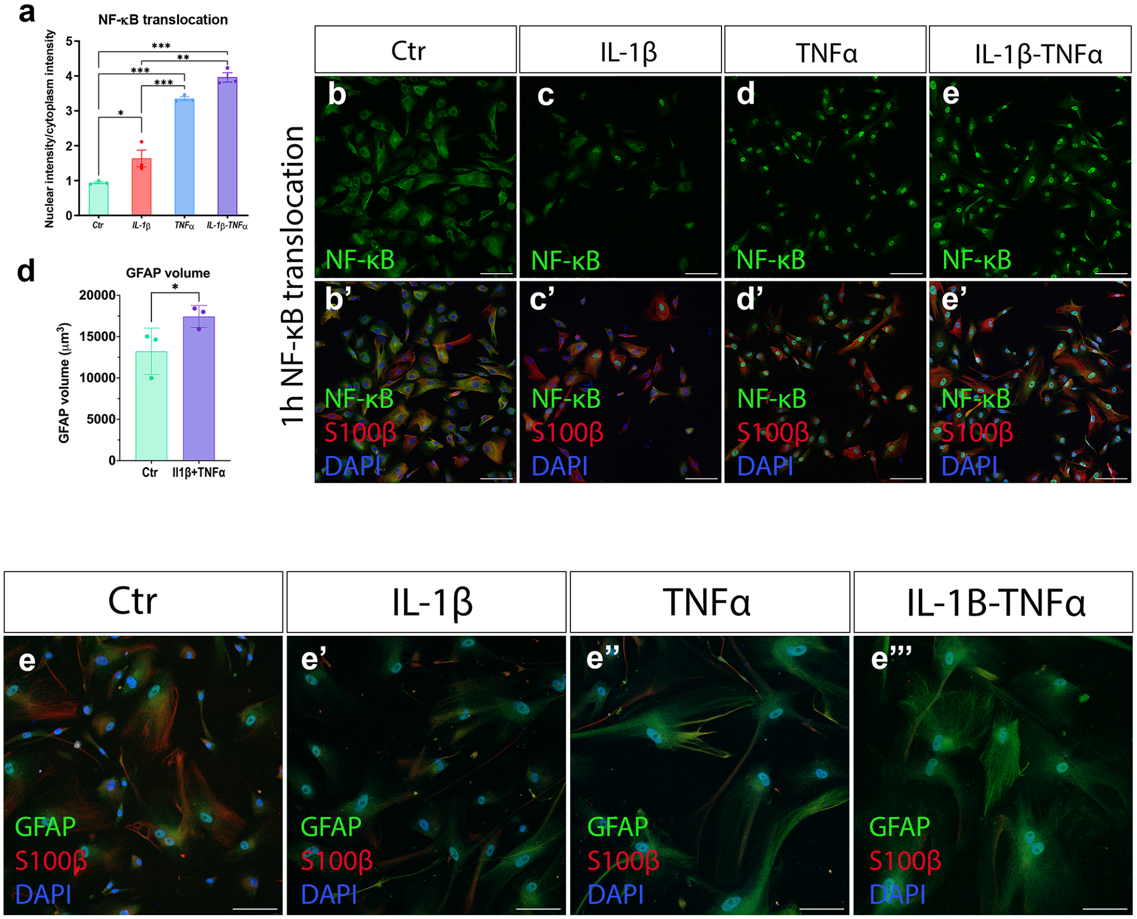


**Suppl. Fig. 3**: **Cytokine treatments and reactivity of hiPSC-derived astrocytes.** (a) Quantification of NF-$\kappa$B nuclear translocation, assessed by the nuclear-to-cytoplasmic intensity ratio in each condition ($n = 3$; $p < 0.05$, $^*p \leq 0.001$, $^{**}p < 0.0001$; one-way ANOVA with Tukey's multiple comparisons test). (b–e') Representative immunofluorescence images showing NF-$\kappa$B (green) localization under different cytokine treatment conditions. (b,b') In control conditions, NF-$\kappa$B remains entirely cytoplasmic. (c,c') In IL-1$\beta$-treated astrocytes, NF-$\kappa$B is partially localized in both the cytoplasm and nucleus. (d,d') In TNF-$\alpha$-treated astrocytes, NF-$\kappa$B is predominantly nuclear. (e,e') In the combined IL-1$\beta$ + TNF-$\alpha$ condition, NF-$\kappa$B is almost entirely nuclear. (b',c',d',e') Merged images showing NF-$\kappa$B (green), S100$\beta$ (red), and DAPI (blue) across conditions. (e–e"') Representative images showing astrocyte morphology at the 7-day timepoint. (e) Control condition, showing resting astrocyte morphology. (e') IL-1$\beta$-treated astrocytes exhibit a slightly enlarged soma. (e") TNF-$\alpha$-treated astrocytes display more pronounced soma enlargement and elongated processes. (e"') IL-1$\beta$ + TNF-$\alpha$-treated astrocytes exhibit the most evident morphological changes, including pronounced cell enlargement and extended processes. Scale bars = 100 $\mu$m. These findings confirm that NF-$\kappa$B nuclear translocation and astrocyte morphological changes are hallmarks of inflammatory reactivity in hiPSC-derived astrocytes.

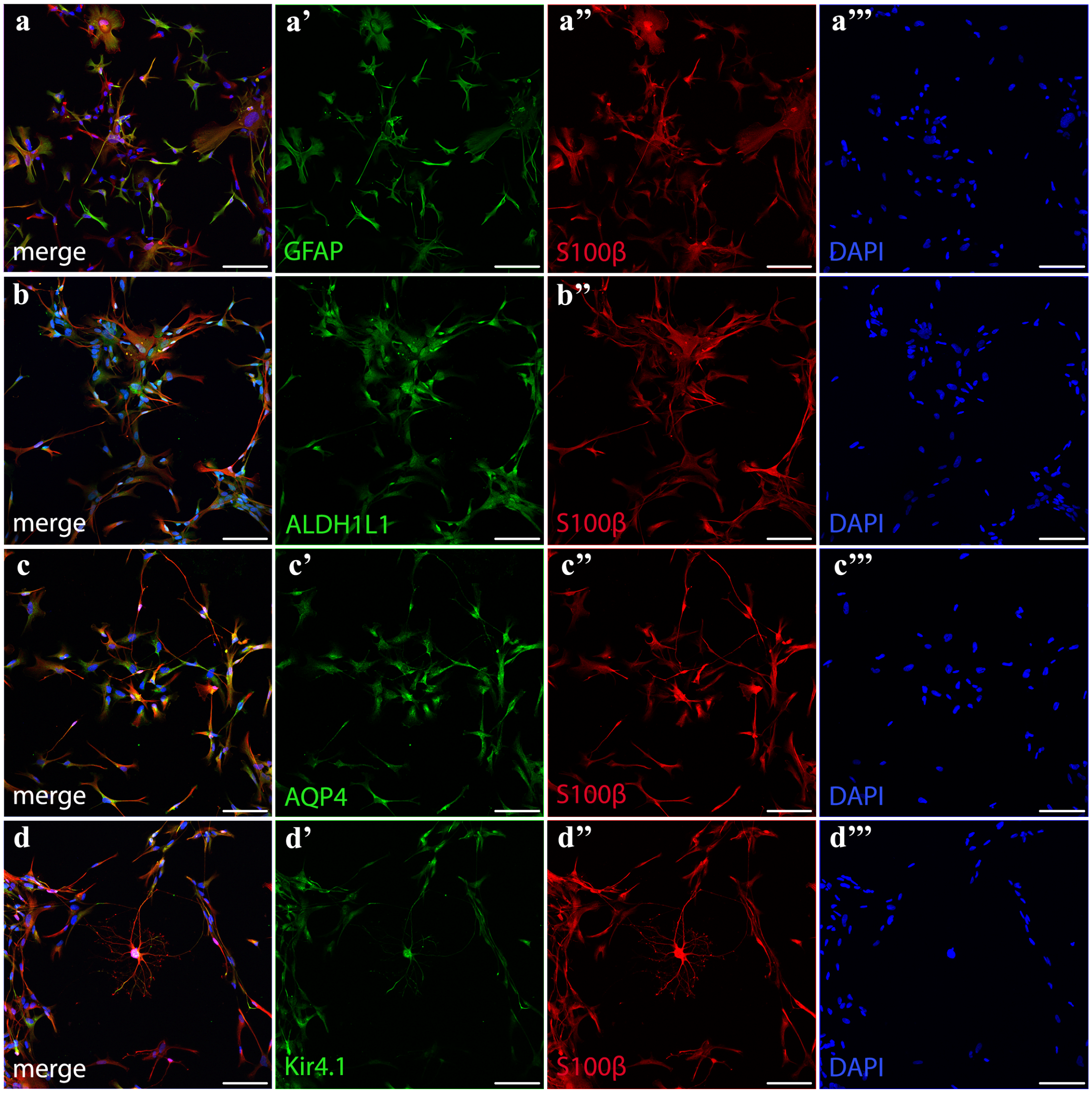


**Suppl. Fig. 4**: **Characterization of hiPSC-derived astrocytes in mixed culture (1).** (a–d) Representative immunofluorescence images showing the expression of astrocytic markers in human induced pluripotent stem cell (hiPSC)-derived astrocytes within mixed cultures. (a) GFAP (green), (b) ALDH1L1 (green), (c) AQP4 (green), and (d) Kir4.1 (green) are co-stained with the astrocytic marker S100$\beta$ (red) and nuclear marker DAPI (blue). Individual channels highlighting the expression pattern of GFAP (a'), ALDH1L1 (b'), AQP4 (c'), and Kir4.1 (d'). GFAP is more pronouncedly expressed in the cytoplasm, ALDH1L1 is present in the astrocytic cytoplasm, AQP4 shows increased expression in both the cytoplasm and membrane, and Kir4.1 is distinctly localized in the astrocytic cytoplasm and membrane. (a",b",c",d") S100$\beta$ (red) effectively delineates astrocyte morphology, appearing more defined compared to hiPSC-derived immature astrocytes. (a"',b"',c"',d"') DAPI (blue) stains cell nuclei. Scale bars = 100 $\mu$m. These findings confirm the astrocytic identity of hiPSC-derived astrocytes in mixed cultures.

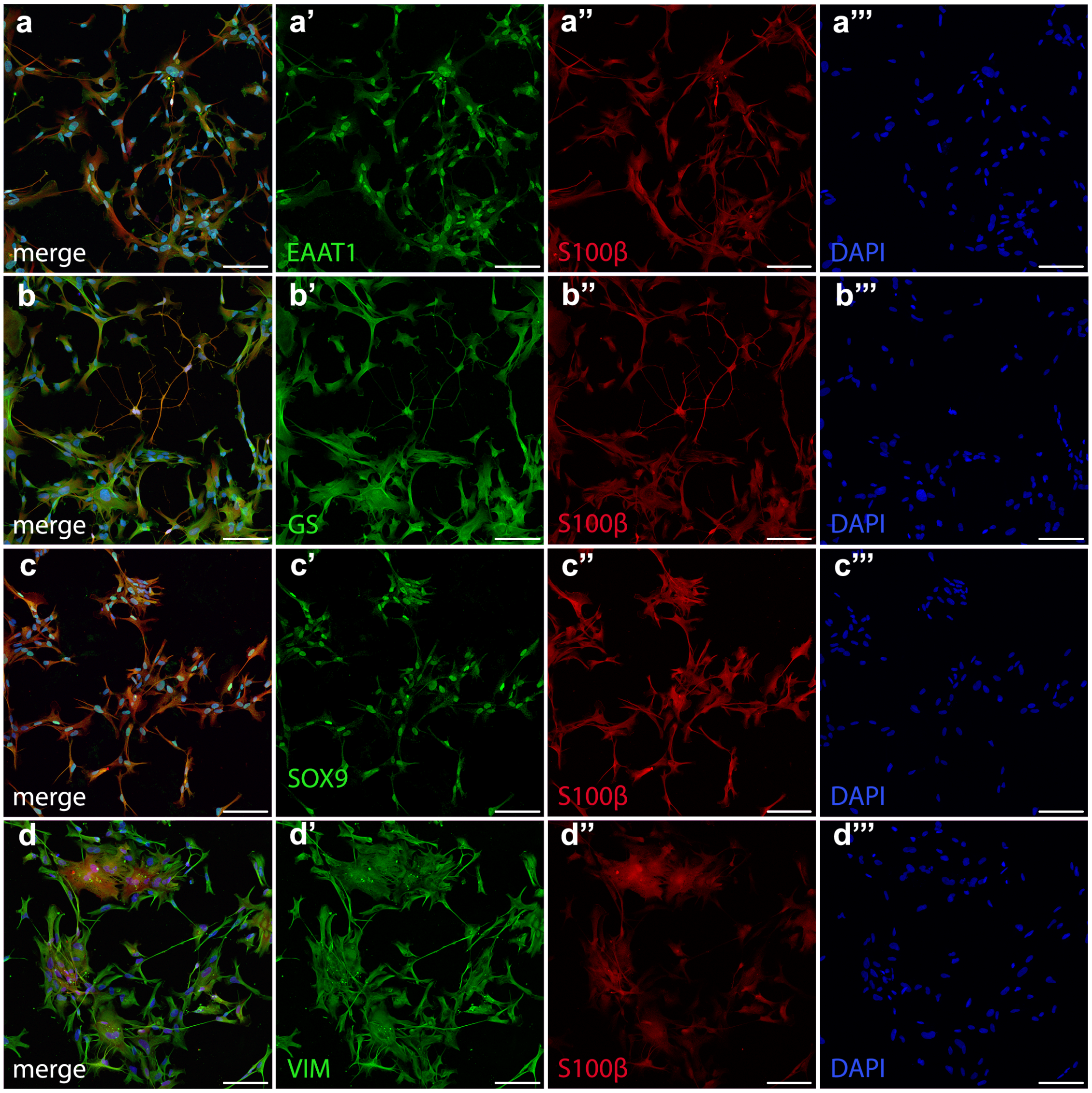


**Suppl. Fig. 5**: **Characterization of hiPSC-derived astrocytes in mixed culture (2).** (a–d) Representative immunofluorescence images showing the expression of astrocytic markers in human induced pluripotent stem cell (hiPSC)-derived astrocytes within mixed cultures. (a) EAAT1 (green), (b) GS (green), (c) SOX9 (green), and (d) VIM (green) are co-stained with the astrocytic marker S100$\beta$ (red) and nuclear marker DAPI (blue). Individual channels highlighting the expression pattern of EAAT1 (a’), GS (b’), SOX9 (c’), and VIM (d’). EAAT1 is localized in both nuclei and cell processes, GS is strongly expressed in the cytoplasm, SOX9 is detected in both cytoplasmic and nuclear compartments with more pronounced nuclear staining, and VIM outlines the morphology of hiPSC-derived astrocytes. (a”,b”,c”,d”) S100$\beta$ (red) effectively delineates astrocyte morphology. (a”’,b”’,c”’,d”’) DAPI (blue) stains cell nuclei. Scale bars = 100 $\mu$m. These findings confirm the astrocytic identity of hiPSC-derived astrocytes in mixed cultures.

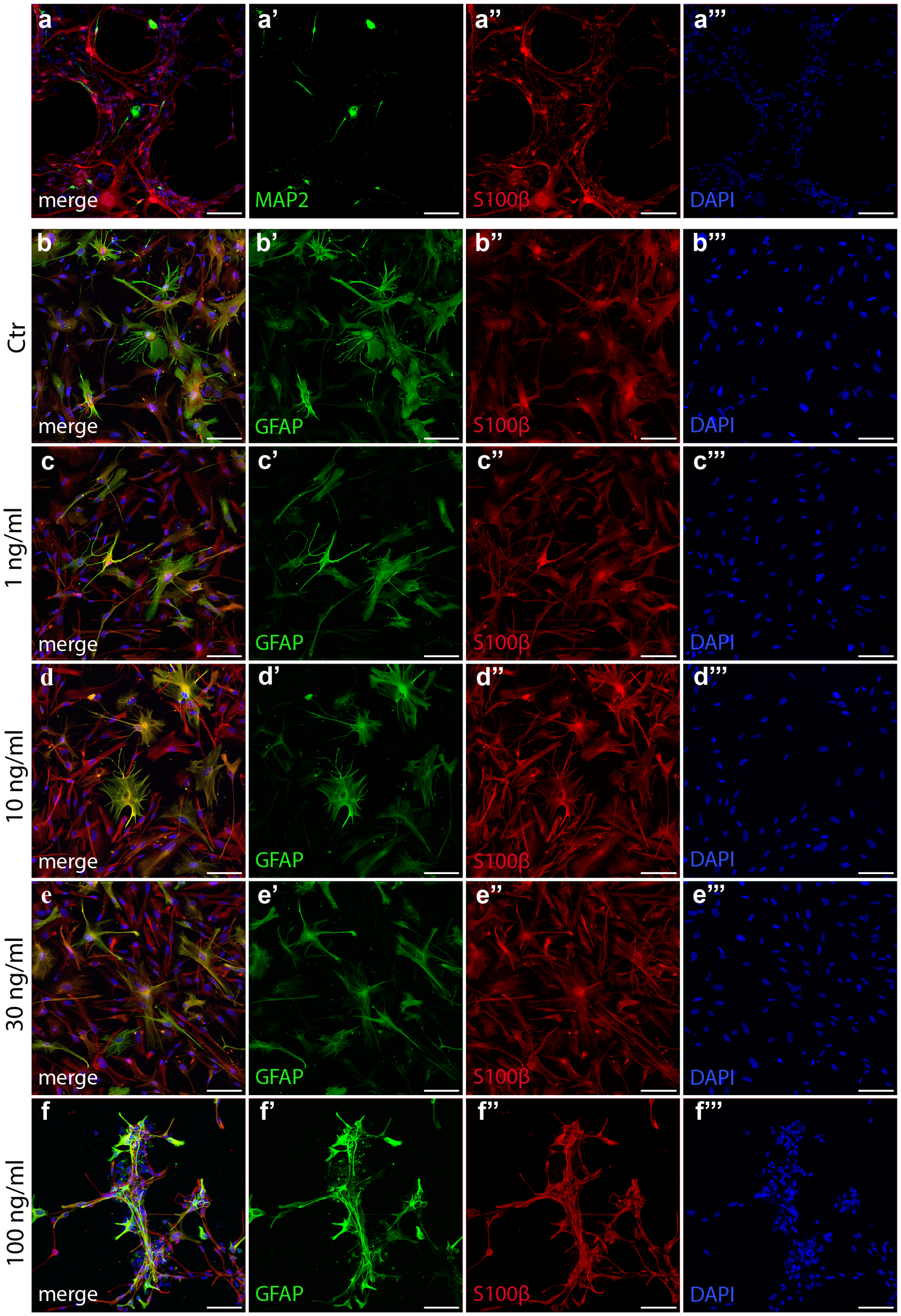


**Suppl. Fig. 6**: **Cytokine treatments and reactivity of hiPSC-derived astrocytes in mixed culture.** (a) Representative immunofluorescence image showing the presence of neurons in mixed cultures, confirmed by staining for MAP2 (green), S100$\beta$ (red), and DAPI (blue). Individual channels highlighting MAP2 (a'), S100$\beta$ (a"), and DAPI (a"'). (b–f) Representative immunofluorescence images showing astrocyte morphology under different cytokine treatment conditions. (b) Control condition, with merged staining for GFAP (green), S100$\beta$ (red), and DAPI (blue). Individual channels showing GFAP (b'), S100$\beta$ (b"), and DAPI (b"') in the control condition. (c–f) Cytokine treatments with increasing concentrations of IL-1$\beta$ + TNF-$\alpha$ (1 ng/mL to 100 ng/mL), showing progressive astrocyte reactivity. (c–c"') 1 ng/mL treatment, (d–d"') 10 ng/mL treatment, (e–e"') 30 ng/mL treatment, (f–f"') 100 ng/mL treatment. GFAP staining (c'–f') highlights astrocyte morphology changes, S100$\beta$ (c"–f") marks astrocytes, and DAPI (c"'–f"') stains nuclei. Scale bars = 100 $\mu$m. These findings confirm astrocyte reactivity and morphological changes in response to increasing cytokine concentrations in mixed cultures.

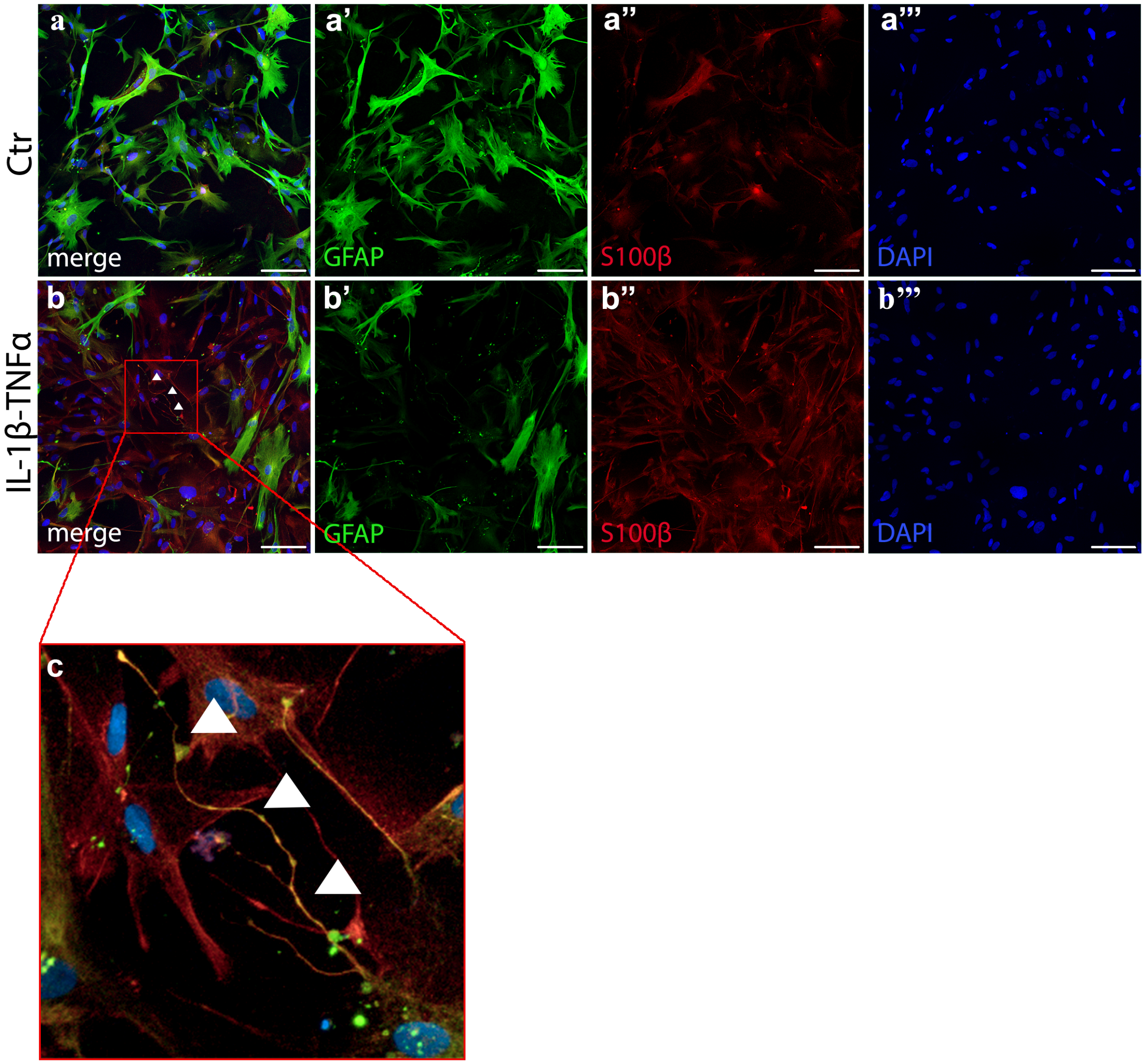


**Suppl. Fig. 7**: **Varicose-projection astrocytes in treated human astrocytes in mixed culture.** (a) Representative immunofluorescence image of astrocytes in the control condition, with merged staining for GFAP (green), S100$\beta$ (red), and DAPI (blue). Individual channels highlighting GFAP (a'), S100$\beta$ (a"), and DAPI (a"') in control astrocytes. (b) Astrocytes treated with IL-1$\beta$ (10 ng/mL) + TNF-$\alpha$ (10 ng/mL) for 7 days, showing the presence of varicose projections (arrowheads). Merged staining for GFAP (green), S100$\beta$ (red), and DAPI (blue). Individual channels highlighting GFAP (b'), S100$\beta$ (b"), and DAPI (b"') in treated astrocytes, with white arrowheads indicating varicose projections. (c) Higher magnification of a varicose projection, with arrowheads marking its structure. Scale bars = 100 $\mu$m. These findings confirm the presence of varicose-projection astrocytes in cytokine-treated human astrocytes within mixed cultures.

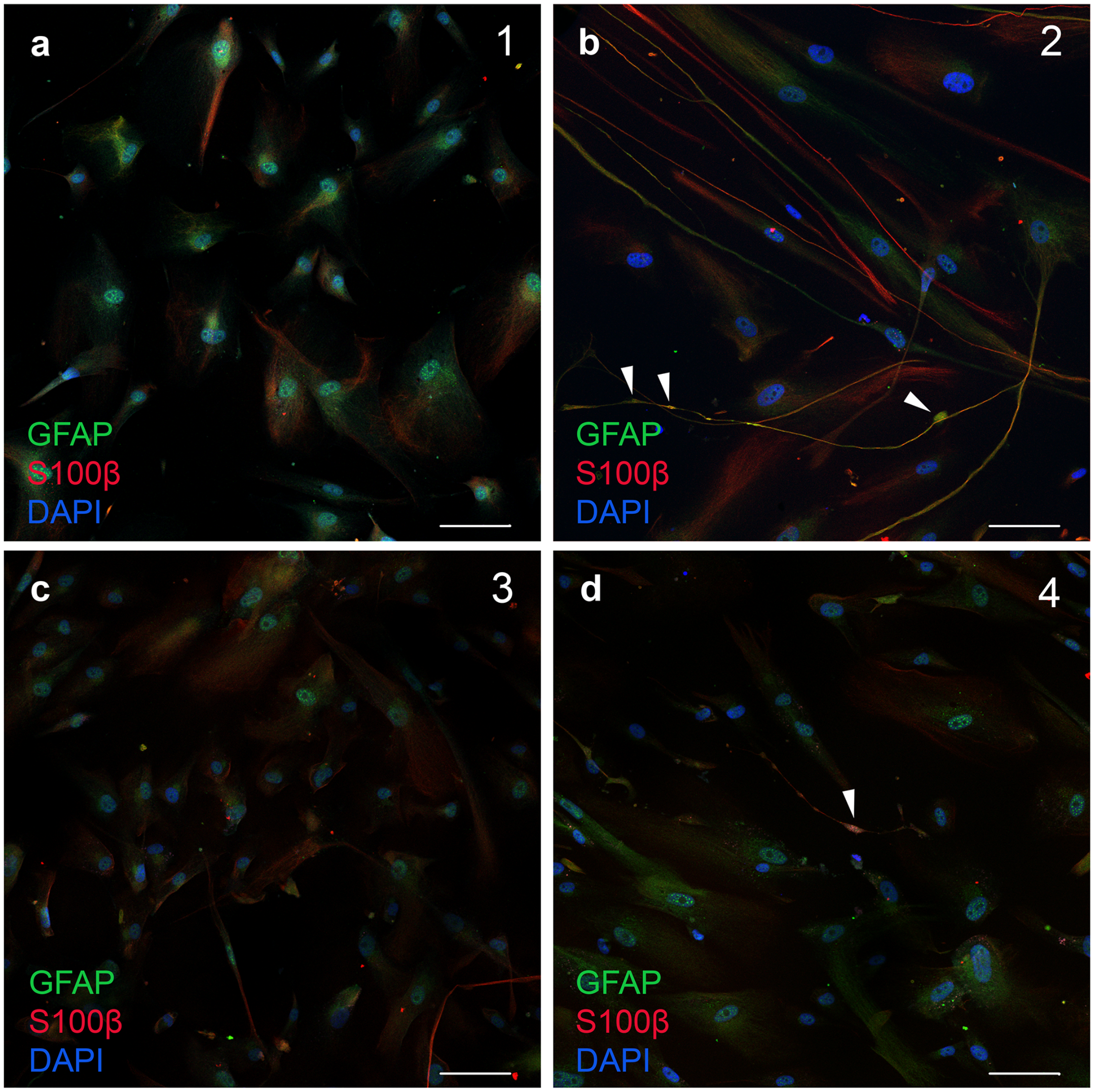


**Suppl. Fig. 8**: **VP astrocyte density increase is reversible one week after removing the cytokine exposure.** (a–d) Representative immunofluorescence images of astrocytes stained for GFAP (green), S100$\beta$ (red), and DAPI (blue) under different conditions. (a) Control condition at 7 days showing an absence of VP astrocytes. (b) IL-1$\beta$ + TNF-$\alpha$ treatment at 7 days, showing a marked increase in VP astrocytes (c) Control at 14 days, confirming no spontaneous VP astrocyte formation over time. (d) Cytokine withdrawal at 14 days, demonstrating a strong reduction in VP astrocytes compared to the cytokine-treated condition at 7 days. White arrowheads point to varicosities. Scale bars = 50 $\mu$m.

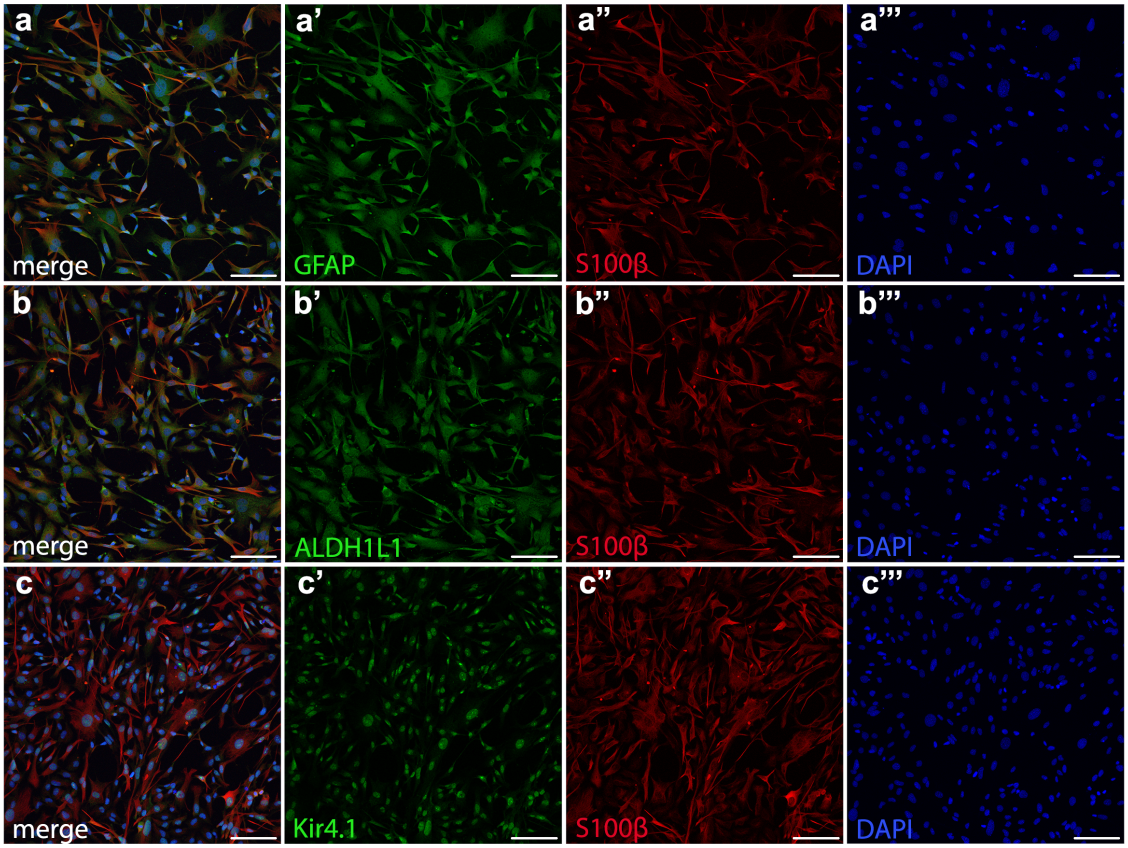


**Suppl. Fig. 9**: **Characterization of ESCs-derived mouse astrocytes 1.** (a–c) Representative immunofluorescence images showing the expression of key astrocytic markers in embryonic stem cell (ESC)-derived mouse astrocytes. (a) GFAP (green), (b) ALDH1L1 (green), and (c) Kir4.1 (green) are co-stained with the astrocytic marker S100$\beta$ (red) and nuclear marker DAPI (blue). Individual channels highlighting the expression pattern of GFAP (a'), ALDH1L1 (b'), and Kir4.1 (c'). GFAP is localized in the cytoplasm, ALDH1L1 is expressed in the astrocytic cytoplasm, and Kir4.1 is predominantly found in the nuclear region. (a",b",c") S100$\beta$ (red) effectively delineates the morphology of ESC-derived astrocytes. (a"',b"',c"') DAPI (blue) stains cell nuclei. Scale bars = 100 $\mu$m. These findings confirm the astrocytic identity of ESC-derived mouse astrocytes based on the expression of multiple astrocytic markers.

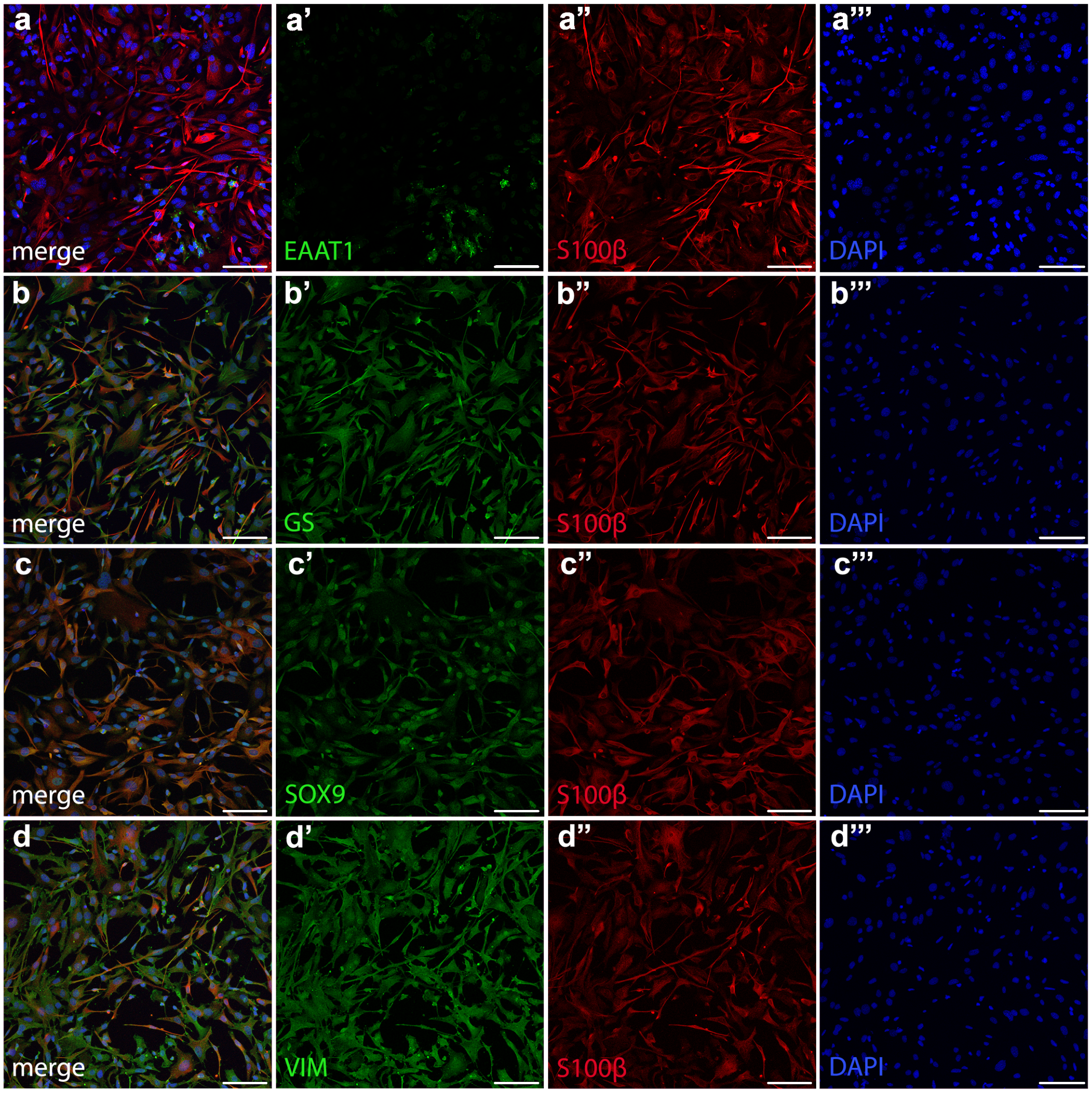


**Suppl. Fig. 10**: **Characterization of ESCs-derived mouse astrocytes 2.** (a–d) Representative immunofluorescence images showing the expression of additional astrocytic markers in embryonic stem cell (ESC)-derived mouse astrocytes. (a) EAAT1 (green), (b) GS (green), (c) SOX9 (green), and (d) VIM (green) are co-stained with the astrocytic marker S100$\beta$ (red) and nuclear marker DAPI (blue). Individual channels highlighting the expression pattern of EAAT1 (a'), GS (b'), SOX9 (c'), and VIM (d'). EAAT1 is predominantly localized in the nucleus with faint expression in cell processes, GS is strongly expressed in the cytoplasm, SOX9 is detected in both the nucleus and cytoplasm, and VIM delineates the morphology of ESC-derived astrocytes. (a",b",c",d") S100$\beta$ (red) effectively marks the morphological features of ESC-derived astrocytes. (a"',b"',c"',d"') DAPI (blue) stains cell nuclei. Scale bars = 100 $\mu$m. These findings further confirm the astrocytic identity of ESC-derived mouse astrocytes based on the expression of multiple astrocytic markers.

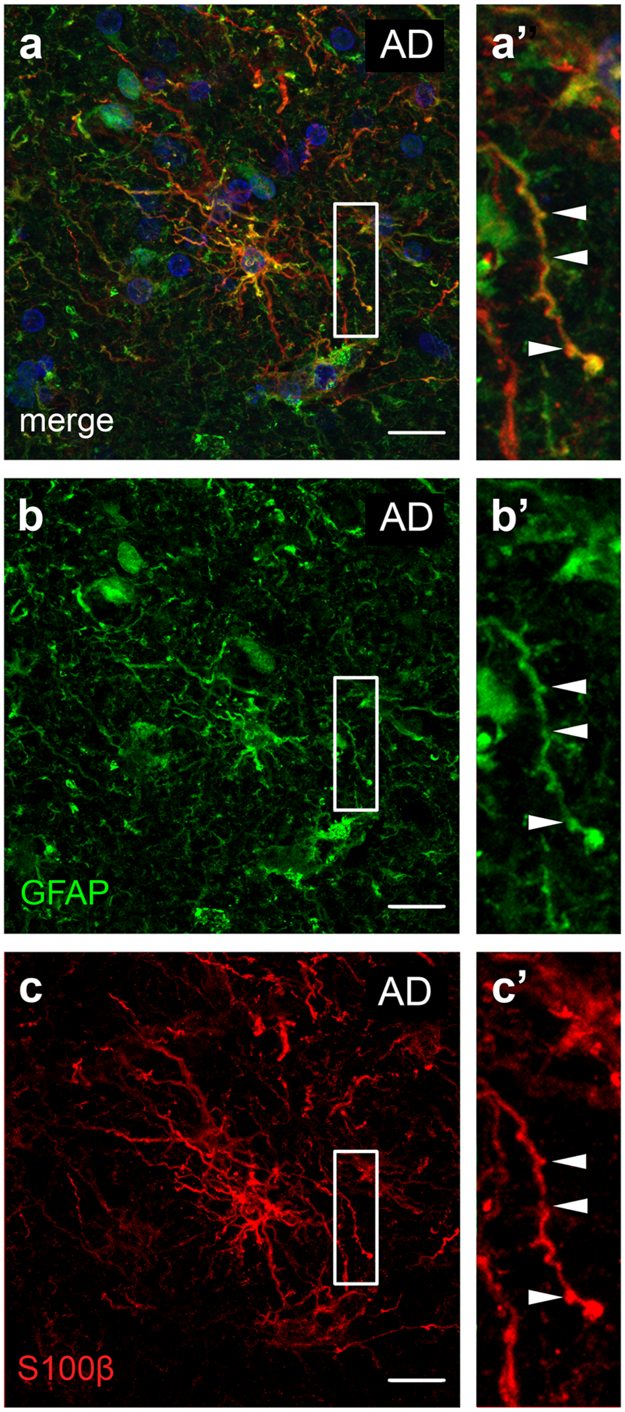


**Suppl. Fig. 11**: **VP astrocytes express both GFAP and S100$\beta$.** Representative immunofluorescence images showing VP astrocytes stained with GFAP (green), S100$\beta$ (red) and DAPI (blue). (a) merge, (b) GFAP only, (c) S100$\beta$ only. (a', b', c') Higher magnification of the boxed regions in (a), (b), and (c), respectively. Arrowheads point to varicosities. Scale bars = 30 $\mu$m.
